# Comprehensive Survey of Conserved RNA Secondary Structures in Full-Genome Alignment of Hepatitis C Virus

**DOI:** 10.1101/2023.11.15.567179

**Authors:** Sandra Triebel, Kevin Lamkiewicz, Nancy Ontiveros, Blake Sweeney, Peter F. Stadler, Anton I. Petrov, Michael Niepmann, Manja Marz

## Abstract

Hepatitis C virus (HCV) is a plus-stranded RNA virus that often chronically infects liver hepatocytes and causes liver cirrhosis and cancer. These viruses replicate their genomes employing error-prone replicases. Thereby, they routinely generate a large “cloud” of RNA genomes which - by trial and error - comprehensively explore the sequence space available for functional RNA genomes that maintain the ability for efficient replication and immune escape. In this context, it is important to identify which RNA secondary structures in the sequence space of the HCV genome are conserved, likely due to functional requirements.

Here, we provide the first genome-wide multiple sequence alignment (MSA) with the prediction of RNA secondary structures throughout all representative full-length HCV genomes. We selected 57 representative genomes by clustering all complete HCV genomes from the BV-BRC database based on k-mer distributions and dimension reduction and adding RefSeq sequences. We include annotations of previously recognized features for easy comparison to other studies.

Our results indicate that mainly the core coding region, the C-terminal NS5A region, and the NS5B region contain secondary structure elements that are conserved beyond coding sequence requirements, indicating functionality on the RNA level. In contrast, the genome regions in between contain less highly conserved structures. The results provide a complete description of all conserved RNA secondary structures and make clear that functionally important RNA secondary structures are present in certain HCV genome regions but are largely absent from other regions. Full-genome alignments of all branches of *Hepacivirus C* are provided in the supplement.

## Introduction

Hepatitis C virus (HCV) is a major public health concern that affects an estimated 71 million people globally [1] causing liver cirrhosis and hepatocellular carcinoma. The virus is a member of the species *Hepacivirus C* and the *Flaviviridae* family, which also includes Dengue, Zika, and yellow fever viruses. HCV is primarily transmitted through blood-to-blood contact, with injection drug use being the most common mode of transmission. It can also be transmitted through unsafe medical procedures, blood transfusions, and from mother to child during childbirth.

HCV is a single-stranded positive-sense RNA virus. The HCV genome is approximately 9.6 kb in length and organized into a single open reading frame (ORF) flanked by two untranslated regions (UTRs) at the 5’ and 3’ ends. The ORF encodes a single polyprotein precursor that is co- and post-translationally cleaved by host and viral proteases into structural (C, E1, E2, p7) and non-structural proteins (NS2, NS3, NS4A, NS4B, NS5A, NS5B), which play critical roles in viral replication and pathogenesis [2, 3, 4]. The HCV genome is known to form several RNA secondary structures [5, 6, 7, 8, 9, 10, 11, 12, 13]. The 5’ UTR contains four structural domains (I-IV). Three of those (II-IV) form a highly structured RNA region called the internal ribosome entry site (IRES), which is responsible for controlling the translation of the viral polyprotein. The IRES allows HCV to circumvent the host cell’s cap-dependent translation initiation mechanism, which is commonly used by cellular mRNAs. The region downstream of SL I including alternative SL II structures [14] contains binding sites for microRNA-122 (miR-122) involved in translation, replication, and RNA stability [7, 15, 16, 17, 18]. The 3’ UTR contains several RNA structures that are involved in regulating viral RNA replication and translation, including a hypervariable region (HVR), a poly-U/UC tract, and a highly conserved RNA secondary structure called X-tail. The X-tail is a conserved RNA stem-loop structure that serves as a binding site for host proteins and is involved in viral replication and translation [7, 11]. Apart from the structural motifs located in the UTRs, HCV showcases RNA secondary structures within its coding region, such as the *cis*-replication element [5, 6, 7, 8, 13, 19, 20].

The HCV genome is highly variable, with eight genotypes and 93 subtypes identified to date, and a sequence diversity of approximately 30 % between genotypes [21, 22], see Figure S1. Its genome contains several complex RNA structures that are critical for viral replication and pathogenesis. Thus, understanding the structure and function of these RNA elements is crucial for the development of effective treatments for HCV infection.

Previous studies have predicted conserved RNA secondary structures in important parts of the HCV genome by different approaches. These studies either analyzed mainly the 5’ and 3’ UTRs as well as the end of the NS5B coding region [7, 13], or they focused on certain HCV genome hotspot regions in the coding regions using covariance analysis with genotype 2 sequences and extending this to other genotypes [8]. Another study confined the prediction of RNA secondary structures in a full-length genome on the genotype 2a JFH-1 isolate, but no other genotypes [6]. The above studies provided important information on conserved RNA secondary structures and their functions in certain HCV RNA genome regions. However, up to the present, a complete survey of all conserved RNA secondary structures in the full length HCV across all genotypes is missing. Therefore, we filtered and manually refined a set of 2 549 HCV full-length genome sequences that fully represent the phylogenetic diversity of all known HCV isolates. From this set, 57 HCV RNA genome sequences were selected that represent the complete phylogenetic tree’s sequence space of HCV genomes.

The *in silico* calculation of full-genome alignments for viral sequences, coupled with the prediction of RNA secondary structures, presents a multifaceted challenge in computational biology. Viral genomes exhibit high genetic diversity, characterized by rapid mutation rates and the presence of insertions and deletions. The complexity for the construction of a multiple sequence-secondary structure alignment (MSSSA) lays at *O*(*m·n*^6^) and is therefore, not computationally feasible. For current construction of MSSSAs, the genomes have to be divided into smaller subsequences for a reliable prediction. Addressing these challenges is pivotal, as such alignments provide crucial insights into the molecular evolution of viruses and their structural-functional relationships. In this context, the development of robust computational methodologies is essential to advance our understanding of viral biology and host-virus interactions.

In this study, we present a full-genome alignment of HCV coupled with RNA secondary structure annotation. The alignment was generated using a semi-automated approach and underwent curation led by experts in the field of HCV and its associated structural elements. Beyond established structures, our study reveals previously unrecognized RNA secondary structures, predicted through computational methods, that exhibit conservation across HCV genomes. Moreover, the results of our alignment – using sequences covering the complete phylogenetic sequence space of HCV isolates – suggest that our predictions likely cover virtually all possible conserved RNA secondary structures.

## Material and Methods

### Data

We downloaded 2,606 HCV genomes (June 01, 2023) from the `BV-BRC` database [23]. To ensure the quality of the data, we filtered the genome status “complete” and excluded the host group “lab”. Notably, despite the genome completeness filter, the majority of entries of this data set (80.5 %) contain incomplete genomes, lacking the UTRs. Among these, 20 genomes were excluded from the analysis due to their sequences containing 10 % or more ‘`N`’s. After identifying duplicated genomes, the data set was refined to a total of 2 549 genomes. Both the original data set and the pre-filtered data set are included in the supplementary files F1 and F2 in `Fasta (.fasta)` format.

### Finding representative genomes

We performed clustering of the pre-filtered data set (2 549 genomes) based on k-mers to select sequences representing the data set. After calculating the k-mer profiles of the input sequences, we performed a dimension reduction by principal component analysis (PCA) followed by clustering using `HDBSCAN` v0.8.27 [24]. `HDBSCAN` resulted in 36 representative genomes which were selected for further analysis, as this method provided comprehensive coverage of the genome information space. For comparison, we clustered sequences with five algorithms: `cd-hit-est` [25, 26], `MMSeqs2` [27], `sumaclust` [28], `vclust` [29], and `HDBSCAN` [24] (see Table S1). The workflow is implemented in `ViralClust` [30]. Despite all filters applied, partial genomes are present in the data set, and thus, the cluster representatives calculated by `HDBSCAN` did too. We removed six sequences manually from the set of representative genomes because they were too short (less than 1 000 nt: `MK468966, MK468983, MK469005, MK468990, and OM896954`), or were not related to a functional polyprotein (`EU862828`). However, it was necessary to manually enlarge our set of representative genomes to fully display the entire spectrum of HCV samples. Utilizing the phylogenetic tree of the pre-filtered data set, see Figure S1 and supplementary file F3, we added 20 genomes representing outliers or subtrees not covered by the clustering results (black squares in Figure S1). Additionally, for comparison to known strains, we added the `NCBI` RefSeq genomes of HCV (NC 038882, NC 004102, NC 009823, NC 009824, NC 009825, NC 009826, NC 009827, NC 030791) to our final set of representative genomes [31]. We removed one sequence (`OM896952`) because of high redundancy with `NC 009824`. Finally, a total of 57 representative genomes covering all eight genotypes of HCV were selected, see supplementary file F4. About 50 % of the representative HCV genomes (27) contain the UTRs (see supplementary information subsection ‘Genome completeness of representative genomes’). The genome length of the selected genomes ranges from 9,036 nt (`MN164872.1`) to 9,711 nt (`NC 009823.1`).

### Multiple sequence alignment and RNA secondary structure prediction

The 57 representative genomes served as input for alignment construction. We computed an initial multiple sequence alignment (MSA) using `MAFFT` v7.520 [32] to identify highly conserved regions, which served as ‘anchors’ for further steps. Anchors are defined as segments in the MSA, requiring a minimum length of 10 nucleotides, and exhibiting an average Shannon entropy value lower than 0.1. Subsequently, we focused our analysis on the subregions between these anchors, utilizing `LocARNA` v2.0.0 [33]. The subregions were then merged into one MSA, followed by an RNA secondary structure prediction of the full-genome alignment with a window-based approach. These steps are implemented in `VeGETA` using `Python` v3.7.12 [34, 35]. Additionally, we intensively examined and slightly curated the alignment manually. Finally, we added the annotation of conserved RNA secondary structures described in the literature as well as novel ones. Based on the nucleotide alignment, we constructed the protein alignment of the representative genomes.

## Results & Discussion

In this study, we present a comprehensive analysis of all conserved RNA secondary structures that occur in the complete sequence space represented by the phylogenetic tree of all HCV isolates (see Figure S1). Thus, the results are supposed to provide a complete description of RNA secondary structures that are of importance for the viral life cycle.

It is known that functionally important RNA secondary structures not only occur in the untranslated regions of mRNAs but also in the coding regions [36]. A variety of molecular mechanisms can be envisioned to be employed by RNA secondary structure elements. RNA secondary structure elements in the protein-coding region can be used for influencing the translational outcome of a given RNA [37], for example by inducing a ribosome frameshift or termination reinitiation. Specific RNA elements can be used for packaging selectively one RNA while another longer RNA species is excluded from packaging by translational inactivation [38]. A translating ribosome may also displace proteins from an RNA secondary structure element of the RNA and by that induce a kind of “burn after reading” degradation of the RNA [39]. Thus, it is important to identify those RNA secondary structures that have been selected for their function from the available sequence space produced by the error-prone replicases of RNA plus strand viruses.

### Basic statistics of the full-genome alignments

Our nucleotide-based alignment contains 57 representative HCV genomes including all eight genotypes. The alignment spans 9 831 residues and 23 061 gaps, averaging approximately 405 gaps per sequence. Approximately 8.5% of the sequences within our alignment contain non-`ACGU` characters, highlighting sequence variations that may have functional significance. Half of the sequences (28/57) exhibit a full 5’ UTR; four sequences lack the 5’ UTR entirely; 12 sequences are deficient in both stem-loops I (SL I) and II (SL II) in the 5’ UTR, and 13 sequences lack only SL I. For the 3’ UTR, only 11 sequences display a complete X-tail structure; three sequences show partial X-tail formations, all others lack the 3’ UTR. The seed sequence (`ACACUCC`) of the first miR-122 binding site of the 5’ UTR directly downstream of the SL I is present in all 32 sequences covering that region; the second miR-122 binding site (`CACUCC`) directly downstream is present in 37/40 sequences, and one other sequence contains `CGCUCC` which would also allow miR-122 binding by G-U base pairing. In the 3’ UTR, the miR-122 seed sequence `ACACUCC` is contained in 41/44 sequences, in contrast, three of the sequences in genotypes 6 and 8 do not contain this site.

We added additional information to our nucleotide alignment: (1) gene annotations (shown as annotation line `#=GC Annotation` in the stk file); (2) the F/ARFP frameshift (notated with ‘f’ in the Annotation line in the stk file); (3) the RNA secondary structures (including pseudoknots) documented in the literature, along with alternative structural configurations (see Table 1); (4) incorporated *in-silico* predicted novel RNA secondary structures, providing a comprehensive view of potential conformations within the HCV genome.

**Table 1.** Conserved RNA secondary structures (SS) in HCV genomes, along with their corresponding `Rfam` model IDs (if available) [55, 56]. We updated and merged six `Rfam` models of v12 into five `Rfam` models (v14.10); confirmed the conservation of a total of 16 previously predicted RNA secondary structures throughout the phylogenetic tree (available in `Rfam` v14.10); and added further 23 novel conserved RNA families into `Rfam` (bold font). ○ indicates that the existing `Rfam` model was updated with this publication. We named novel structures according to their position in the genome JFH-1 (S — Start; E – End). *** – During the review process of this manuscript, `Rfam` numbers will substituted here.

| RNA SS | Genomic region | Alignment Position |  | JFH-1 Position |  | Rfam v12 | Rfam v14.10 |
| --- | --- | --- | --- | --- | --- | --- | --- |
|  |  | S | E | S | E |  |  |
| SL I | 5' UTR | 13 | 28 | 5 | 19 | RF00061 | RF00061◦ |
| SL II | 5' UTR | 52 | 126 | 43 | 117 | RF00061 | RF00061◦ |
| SL III | 5' UTR | 133 | 334 | 124 | 322 | RF00061 | RF00061◦ |
| SL IV | 5' UTR/C | 346 | 360 | 334 | 348 | RF00061 | RF00061◦ |
| SL V | C | 400 | 435 | 388 | 423 | RF00620 | RF00620◦ |
| SL VI | C | 439 | 519 | 427 | 507 | RF00620 | RF00620◦ |
| **SL 562** | C | 574 | 595 | 562 | 582 |  |  |
| SL 588 | C | 601 | 678 | 588 | 665 |  | RF04220 |
| SL 669 | C | 684 | 761 | 671 | 748 |  | RF04221 |
| J 750 | C | 762 | 839 | 749 | 826 |  | RF04219 |
| SL 833 | C | 846 | 869 | 833 | 856 |  | RF00*** |
| SL 1414 | E1 | 1 430 | 1 467 | 1 414 | 1 451 |  | RF00*** |
| (former SL 1412) |  |  |  |  |  |  |  |
| **SL 1850** | E1 | 1 881 | 2 013 | 1 850 | 1 982 |  |  |
| **SL 2313** | E1 | 2 356 | 2 433 | 2 313 | 2 390 |  |  |
| SL 2531 | E2 | 2 574 | 2 591 | 2 531 | 2 548 |  | RF00*** |
| SL 2549 | E2/p7 | 2 592 | 2 636 | 2 549 | 2 592 |  | RF00*** |
| **SL 3308** | NS2/NS3 | 3 352 | 3 644 | 3 308 | 3 600 |  |  |
| **SL 3844** | NS3 | 3 888 | 3 952 | 3 844 | 3 908 |  |  |
| **SL 4005** | NS3 | 4 049 | 4 099 | 4 005 | 4 056 |  |  |
| **SL 4214** | NS3 | 4 258 | 4 223 | 4 214 | 4 279 |  |  |
| **SL 4527** | NS3 | 4 571 | 4 591 | 4 527 | 4 547 |  |  |
| **SL 4621** | NS3 | 4 665 | 4 725 | 4 621 | 4 681 |  |  |
| **SL 4691** | NS3 | 4 735 | 4 788 | 4 691 | 4 744 |  |  |
| **SL 5016** | NS3 | 5 060 | 5 125 | 5 016 | 5 081 |  |  |
| **SL 5128** | NS3 | 5 172 | 5 249 | 5 128 | 5 205 |  |  |
| **SL 5357** | NS4A | 5 401 | 5 448 | 5 357 | 5 404 |  |  |
| **SL 5647** | NS4B | 5 691 | 5 777 | 5 647 | 5 733 |  |  |
| **SL 6027** | NS4B | 6 071 | 6 197 | 6 027 | 6 153 |  |  |
| **SL 6270** | NS5A | 6 314 | 6 410 | 6 270 | 6 366 |  |  |
| **SL 6371** | NS5A | 6 415 | 6 446 | 6 371 | 6 402 |  |  |
| **SL 6530** | NS5A | 6 574 | 6 689 | 6 530 | 6 645 |  |  |
| **SL 7516** | NS5A | 7 584 | 7 603 | 7 516 | 7 535 |  |  |
| **SL 7536** | NS5A | 7 604 | 7 702 | 7 536 | 7 634 |  |  |
| **SL 7816** | NS5B | 7 893 | 7 905 | 7 816 | 7 828 |  |  |
| J 7880 | NS5B | 7 959 | 8 073 | 7 882 | 7 996 |  | RF00** |
| SL 8001 | NS5B | 8 077 | 8 126 | 8 000 | 8 049 |  | RF00** |
| **SL 8075** | NS5B | 8 152 | 8 174 | 8 075 | 8 097 |  |  |
| **SL 8299** | NS5B | 8 376 | 8 396 | 8 299 | 8 334 |  |  |
| SL 8670 | NS5B | 8 722 | 8 803 | 8 645 | 8 726 |  | RF00*** |
| 5BSL1 | NS5B | 9 194 | 9 231 | 9 117 | 9 154 |  | RF04218 |
| 5BSL2 | NS5B | 9 276 | 9 335 | 9 198 | 9 257 | RF00468 | RF00468◦ |
| 5BSL3.1 | NS5B | 9 361 | 9 406 | 9 283 | 9 328 | RF00260 | RF00260◦ |
| 5BSL3.2 | NS5B | 9 410 | 9 455 | 9 332 | 9 377 | RF00260 | RF00260◦ |
| 5BSL3.3 | NS5B | 9 467 | 9 497 | 9 389 | 9 419 | RF00260 | RF00260◦ |
| SL IV | NS5B | 9 505 | 9 533 | 9 427 | 9 452 | RF00469 | RF00260◦ |
| X-tail | 3' UTR | 9 727 | 9 826 | 9 579 | 9 678 | RF00481 | RF00481◦ |

The protein alignment of the 57 representative genomes encompasses a total of 3 017 residues, with an average of 37 gaps (2 136 gaps in total). A minor proportion of not characterized amino acid characters (0.0015 %) accounting for a total of 262 occurrences, can be found in the alignment. These ‘`X`’s are based on ‘`N`’s in the sequences downloaded from `NCBI`. We provide the nucleotide, protein, and combined alignments in the supplementary material with several formats, such as `Stockholm (stk)`, `ClustalW (aln)`, and `Fasta (fasta)` (see Files F5 and F7), which can be conveniently visualized using tools such as `ClustalX` [40, 41], `Jalview` [42] or `Emacs RALEE` mode [43]. We have visualized the possible color codes available in `Emacs RALEE` mode in Figure S4 based on the examples SL V and SL VI.

### Alignment confirms previously annotated RNA secondary structures

The full-genome alignment of HCV genomes reveals the presence of well-characterized RNA secondary structures, see Table 1, that are consistent with the existing literature [6, 7, 8, 15]. Therefore, the above alignment is validated by its prediction of these structures which had been demonstrated to be functional *cis*-elements involved in the regulation of HCV RNA translation and replication. These known structural elements include the highly conserved IRES (see Figure S2 and S3), which plays a crucial role in HCV translation initiation [7, 15]. Additionally, stem-loop structures within the core region (see Figure S5) and NS5B coding region (see Figure 1 and S8 - S12), such as the *cis*-replication element (CRE, composed of 5BSL3.1, 5BSL3.2, 5BSL3.3) and the highly conserved X-tail in the 3’ UTR (both conformations) (see Figure 2 and S13), were identified. Moreover, we predicted several conserved RNA secondary structures in the coding region of HCV genomes (see Figure 1, S5, S8, S9, S10, S11, and S12) in agreement with the literature [6, 7, 8]. The identification of these known structures in the core coding region [44, 45] and in the NS5B region [6, 8] not only validates the accuracy of our alignment but also underscores their functional importance in HCV biology: their conservation across different HCV genotypes further highlights their critical roles in viral replication, translation, and infection.

**Figure 1:**
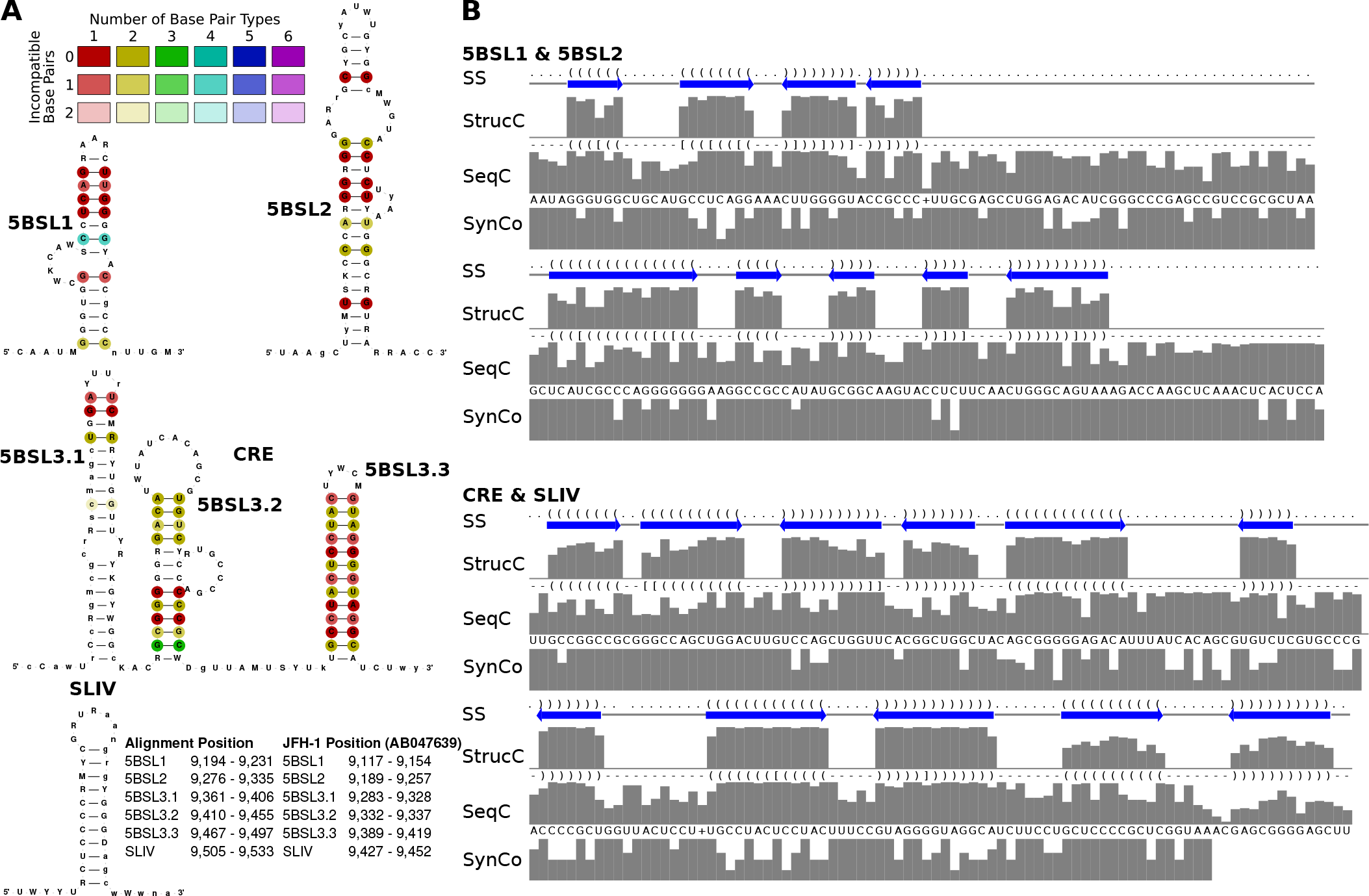
Known RNA secondary structures in the downstream NS5B coding region including the *cis*-replication element (CRE, 5BSL3.2) which are relevant for replication control. (A) RNA secondary structures are colored by the number of base pair types illustrating the extent of covariations in double-stranded regions. The SL IV (or 5BSL3.4) contains the NS5B stop codon in the apical loop. The structure was visualized using `R2DT` [53]. The nucleotide sequence shows the most informative sequence (IUPAC code) calculated by `RNAalifold` based on the alignment. Lowercase letters indicate gaps in the alignment column. (B) RNA secondary structure dot-bracket annotation (SS), structure consensus (StrucC), sequence consensus (SeqC), and the fraction of nucleotides used from synonymous codons (SynCo). Thereby, a low value indicates that only a few nucleotide(s) out of all nucleotides possible for synonymous codons are actually used by the different HCV isolates, indicating a high degree of primary sequence conservation which goes beyond the requirements of the coding sequence. This provides evidence that a conserved functional RNA element may overlap with the coding sequence. Alignments shown in Figure S11 and S12.

**Figure 2:**
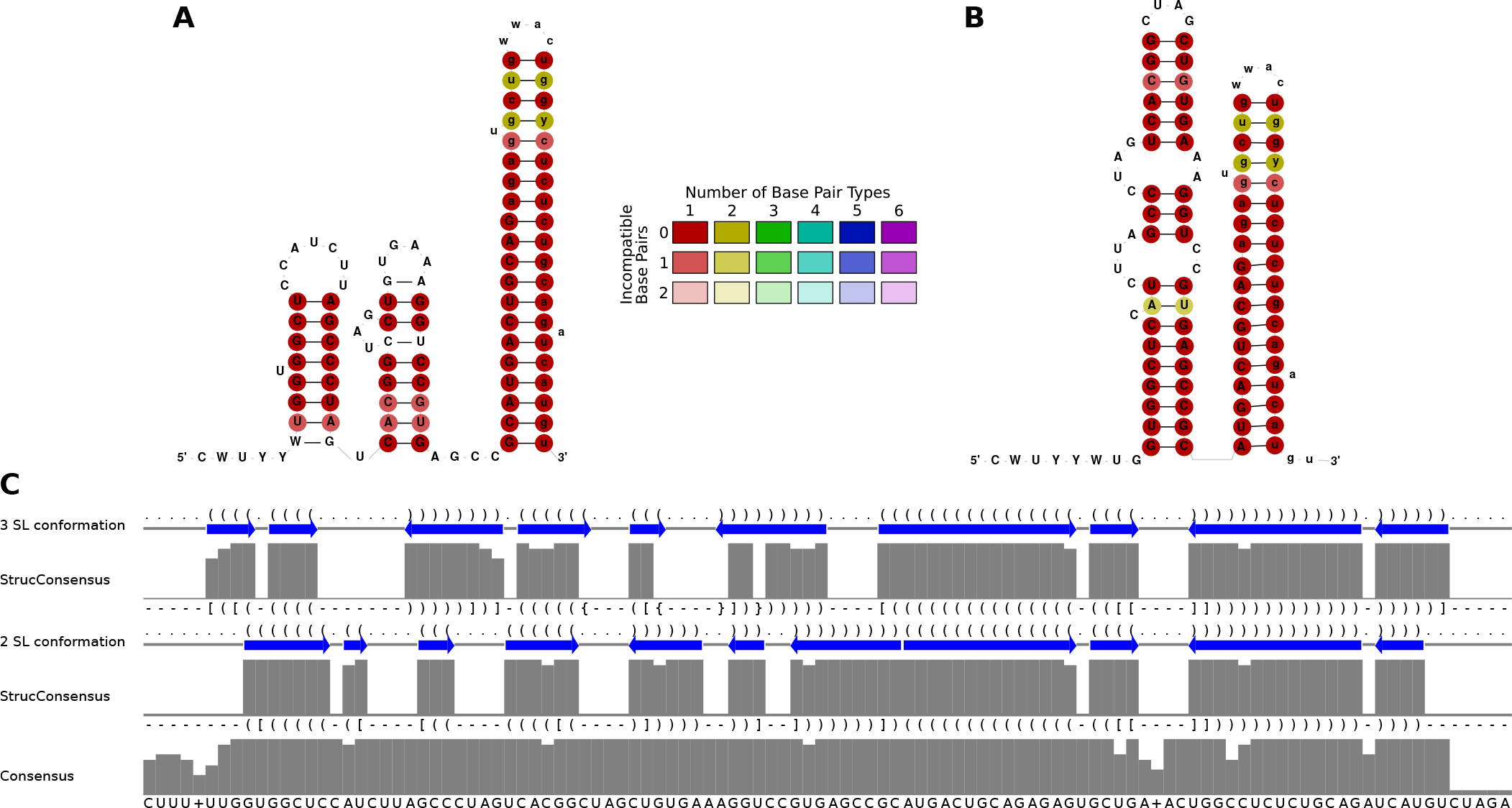
The two alternative structures of the highly conserved 3’ X region of the 3’ UTR. (A) The conformation with the three stem-loops SL 1, 2, and 3. In this form, the apical loop of SL 2 can make a long-range interaction (LRI) with the apical loop of the CRE/5BSL3.2 (“kissing loop” interaction). Among the selected sequences, only 11 isolates had a complete 3’ UTR sequence (please see the additional alignment of only 3’ X sequences in Figure S13 and F9). (B) The conformation with SL 2 and 3 restructured to form the DLS that is speculated to be involved in HCV RNA genome dimerization [50, 51, 54]. (C) Additional sequence and RNA secondary structure features of the consensus of the 11 isolates, with two alternative dot-bracket outputs. As in parts of the 5’ UTR, the strong conservation of the primary sequence indicates that both overlapping structures shown in (A) and (B) may be functionally important, thereby limiting the extent of possible covariations in the RNA secondary structure regions of each conformation.

The CRE/5BSL3.2 and the 5BSL3.3 are important for HCV replication [19, 20]. These functional aspects are reflected by the high conservation of the 5BSL3.2 and 3.3 (see Figure 1 and S12), not only in their RNA secondary structure regions but also in their single-stranded loops and bulges, which are conserved beyond coding sequence requirements, since only selected nucleotides are actually used from those possible in synonymous codons (see Figure 1 and S12). Likely, in the early phase after HCV infection, the CRE apical loop can interact with the SL 2 of the 3’ X region by forming a “kissing loop” interaction [46], and this interaction may stabilize its SL 2, SL 3 conformation [47, 48]. At a time when sufficient amounts of NS5B replicase have been translated, NS5B can bind the CRE/5BSL3.2 and 5BSL3.3 [20, 49], thereby disabling the CRE – SL 2 interaction and allowing formation of the dimerization linkage sequence (DLS) [50, 51]. Binding of NS5B to the CRE then is supposed to be involved in starting RNA minus strand synthesis at the HCV RNA 3’ end, whereas the role of the DLS and its putative role in dimerization of the full-length HCV genome in this process is not yet fully understood. These functional aspects in turn validate our alignment approach for identifying functionally important RNA secondary structures.

Similar constraints may apply to the region including SL II and the preceding sequence between SLs I and II is highly conserved in the primary sequence due to two alternative conformations that fulfill different tasks in the viral life cycle [52]. The classical conformation of SL II allows binding of two molecules miR-122 and has roles in HCV genome replication [16], promoting translation [17] and stabilization of the genome against nucleolytic degradation [18]. The alternative conformation SL II^*alt*^ [14], however, appears to have a role in HCV assembly [52]. Also, the SL IIId and its conserved alternative form SL IIId* which showed up previously [7] in the IRES (see Figure S2 and S3) and is predicted to be slightly more stable than the classical SL IIId. We can only speculate if this alternative SL IIId* represents a structure that may be important in the IRES when not bound to ribosomes.

Additional information about the validated RNA secondary structures, further affirming the quality of our alignment, can be found in the supplementary materials subsection ‘Alignment confirms previously annotated RNA secondary structures – Additional Information’.

### Improvements to Rfam Virus Families

We improved the models in the `Rfam` database [55, 56] (see Table 1) to ensure comprehensive coverage of the entire phylogenetic clade of the HCV sequences. In total, we identified 39 conserved structural regions across the HCV genomes, of which 23 were novel. We updated the six `Rfam` families from release 12 to nine families in release 14.10, of which all are validated by the literature. The HCV Rfam families were reviewed for covariance support with `R-scape` [57]. Only a small number of base pairs (1 to 3) exhibited covariance support in each family, and this consistency was observed across families of both non-coding and coding sequences/regions. The remaining 30 structured regions will be used to create additional `Rfam` families in future releases. In the following, we compare the five models to the previously well-described models from `Rfam` v12: (1) We reduced the IRES model (`RF00061`) from 79 to 51 sequences, spanning 356 nucleotide positions in the alignment (previously 413). The new model includes now SL I (from 30 HCV genomes) and SL II (covered by 41 genomes), which was absent due to sequencing problems of the very 5’ genome end in previous times. Importantly, the new model includes now SL IV from 51 HCV genomes, which is located at the transition from 5’ UTR to core gene, containing the start codon of the polyprotein. (2) The SL V and SL VI model (`RF00620`) was updated from 36 sequences to 56 (`MK548369` excluded because of non-ACGU characters) and now spans an alignment length of 136 positions. The previous model contained 153 alignment positions, indicating a major reduction of gaps in the novel alignment. SL V was reduced by one base pair and SL VI by three base pairs. (3) The 5BSL2 model (`RF00468`) has now been reduced from 110 to 57 genomes. This measurement allows us to not compose a bias towards closely related, highly over-represented sequences. Our alignment expands the stem-loop by four base pairs. (4) The CRE model (`RF00260`) is now represented by all 57 selected genomes (previously 52). Only 5BSL3.2 of the CRE has been included in the old model, therefore the new model spans now 183 nucleotide positions (instead of 51 nucleotides) including the complete CRE (5BSL3.1, 5BSL3.2, and 5BSL3.3) and SL IV. (5) The model of SL IV (`RF00469`), located at the transition from NS5B gene to 3’ UTR, containing the stop codon of the polyprotein, comprised 110 sequences. This structure is now naturally merged into model `RF00260`. (6) Lastly, the X-tail model (`RF00481`) now contains only 11 sequences, the old model contained 22 HCV genomes. However, the old model only contained sequences of genotype 1 – 3. Therefore, although the total number of sequences has been reduced, the variety of the X-tail has been enlarged by our new model and is now spanning genotypes 1 – 6. We expanded SL I by one base pair. In total, we were able to confirm 16 structural elements throughout the entire phylogenetic tree of HCV with were previously confirmed [7, 8, 13], see Figure 1, 2, S2, S5, S6, S8. We added these and the 23 novel HCV RNA secondary structures to `Rfam` v14, see Figure 3 and S14. An information page for all HCV models in `Rfam` is provided at the following link: https://rfam.org/viruses/hcv

**Figure 3:**
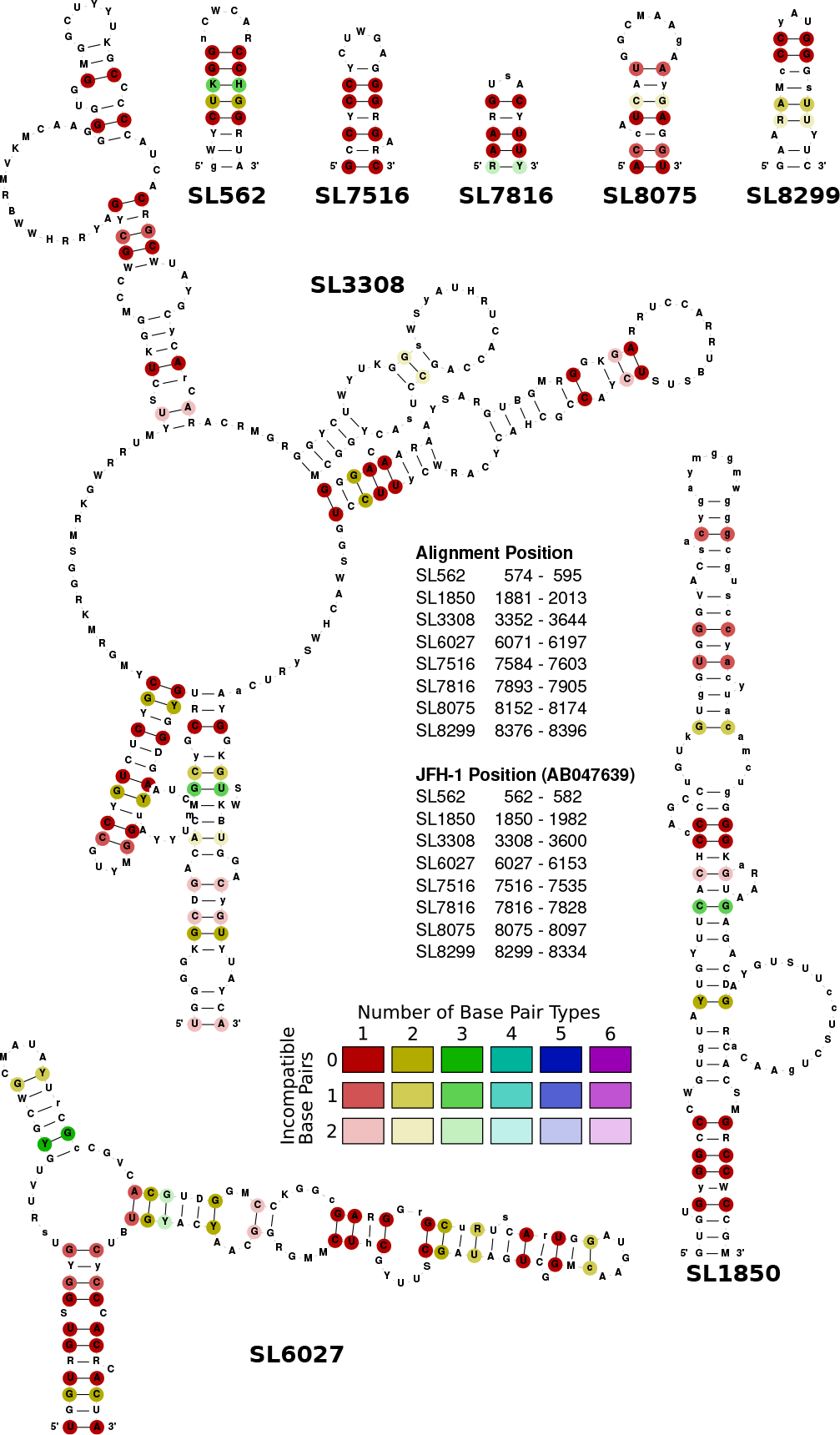
Eight novel selected conserved RNA secondary structure candidates from coding regions of the HCV alignment. RNA secondary structures are colored as in Figure 1 A.

### New RNA secondary structure element candidates

Our analysis revealed several novel RNA secondary structures in the coding region using our *in silico* prediction method (see Figure 3 and S14). We selected novel RNA secondary structures based on their (i) conservation at the sequence and structural level in the representative sequences regarding compensatory mutations and (ii) the use of synonymous codons especially in the hairpin region. We predicted 23 novel candidates to likely be functional elements due to their RNA secondary structure conservation which shows compensatory mutations despite being placed in the coding region, including the restricted use of synonymous codons (Figure 3 and S14). SL 562 (see Figure 3) is one example for a novel predicted short hairpin, which shows compensatory mutations despite being placed in the coding region of the core protein. SL 562 is represented in all 57 sequences highly conserved and structured.

At alignment pos. 7 582 (SL7516, see Figure 3; and JFH-1 pos. 7 516), an RNA secondary structure element with a seven base pair stem and a five nucleotide loop is predicted to be conserved that shows good conservation in the `StructConsensus`, `Consensus` as well as in the low number of exchanges in synonymous codons, indicating conservation beyond coding sequence requirements (see Figure 3 and S15). In these terms, this newly predicted element is better conserved than the J 7880 element which was shown to be functional in early HCV replication [6], suggesting that this new element may have functional importance, even though the degree of its conservation does not fully reach that of the CRE. Similarly, newly predicted RNA secondary structures at positions 7 893 (SL 7816, 7 816 in JFH-1), pos. 8 152 (SL 8075, 8 075 in JFH-1, and pos. 8 376 (SL 8299, 8 299 in JFH-1) are well conserved and are good candidates for functional RNA elements (see Figure 3 and S15). In contrast, some presumable RNA structures in the E1/E2 region, located at alignment positions 1 430, 2 574, and 2 592 in the alignment (SL 1414, SL 2531, and SL 2549; JFH-1 pos. 1 414, 2 531, 2 549; see Figure S6 and S7), are not well enough conserved in terms of StructConsensus (StrucC), Consensus sequence (SeqC) and the limited use of synonymous codons (SynCo) to suggest a possible function.

The above predicted well-conserved structures represent previously unrecognized RNA elements conserved in the representative HCV genomes (see Figure 3 and S14 - S16), highlighting the complexity and diversity of RNA secondary structures within this viral species. The discovery of these novel structures opens up new avenues for understanding their potential functional roles in HCV replication, translation, and pathogenesis. Further investigations are warranted to experimentally validate and explore the functional significance of these newly identified RNA secondary structures in the context of HCV biology.

### A detailed sequence and RNA secondary structure comparison reveals hints into incongruent evolution

Consensus structures are defined by base pairs that are conserved despite substitutions in the underlying sequence. In other words, base pairs that structurally correspond to each other are usually formed by pairs of nucleotides that are homologous according to their position in the sequence context. This is not always the case, however, as demonstrated by the example of the 5BSL2 stem-loop structures from three representative HCV isolates, Figure 4. From a coarse-grained perspective, there is a consensus comprising three helical substructures. A more detailed analysis of the individual stem-loop structures, however, not only shows the expected overall conservation of the structure but also surprising differences. In addition to the expected variation, e.g., of the presence or absence of base pairs at the ends of individual helices or variations in loop sizes, we observe that the innermost helix and the hairpin loop are not formed by homologous nucleotides. Instead, a well-conserved stretch of five nucleotides forming the helix in the consensus (green) is shifted by one position in `MW689971` and three positions in `AY232740` and `NC 009827`, each. As a consequence, the terminal hairpin is conserved as a structural feature, but its individual base pairs are formed by different, non-homologous sequence positions. A similar situation is visible in the middle (red) and outer (blue) stem. The nucleotides forming the middle stem are shifted by four nucleotides relative to the consensus.

**Figure 4:**
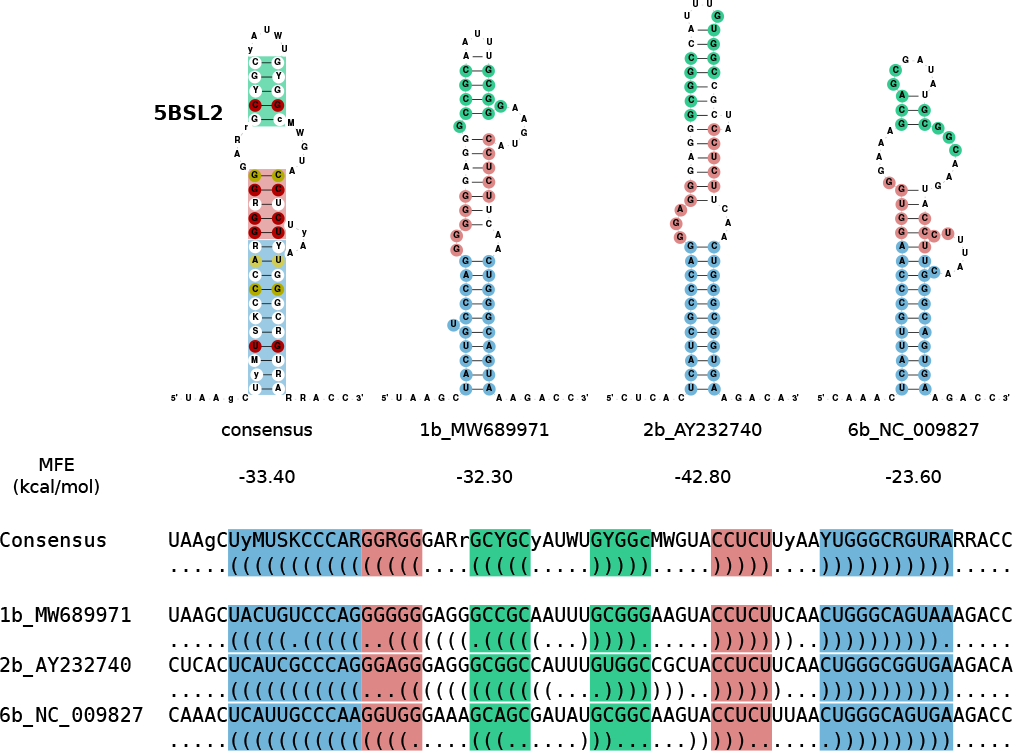
5BSL2 as an example of *incongruent evolution*. The consensus RNA secondary structure differs from the structures into which individual sequences would fold, with sequence and structure shifted relative to each other. A helix of five nucleotides (green) in the consensus is shifted by one position in `MW689971` and three positions in `AY232740` and three positions in `NC 009827`. The nucleotides that form the middle (red) stem are shifted by four nucleotides compared to the consensus.

The conservation of secondary structures realized by non-homologous base pairs was termed *incongruent evolution* [58] in contrast to the more familiar and much more frequent congruent case. Incongruent evolution can be understood as divergence of sequence alignment and structure alignment. Starting from a sequence alignment, such as the one shown on the bottom of Figure 4, this leads to an apparently poor conservation of the structure, indicated by the gray base pairs in the consensus structure. On the other hand, focusing on the structure (shown here by helices in corresponding positions) results in mismatches (indicated by the colored intervals). It has been shown in [59] that incongruences of sequence and structure can be explained mechanistically, e.g. by flexible structural intermediates. Selection pressures act independently, i.e., in different functional contexts, to preserve sequence and secondary structure are particularly conducive to incongruent patterns [36, 60, 61] such as the ones in the 5BSL2 stem-loop structure.

## Conclusion

We presented the first comprehensive genome-wide multiple sequence alignment (MSA), incorporating computational predictions of RNA secondary structures across the entire HCV genomes (see F5 - F9, and Table 1). Our selection of 57 representative genomes across the entire phylogenetic tree is based on clustering all complete HCV genomes from the BV-BRC with HDBSCAN using k-mer distributions and dimension reduction. We added manually the RefSeq sequences. The inclusion of annotations for previously identified features, such as genome annotations, secondary structures, pseudoknots, and alternative structures facilitates seamless comparisons with other research studies. Our in-depth analysis included conservation of the predicted RNA secondary structures, covariance in structured stem regions, and the use of nucleotides in synonymous codons. The latter output not only provides information about the overall conservation of a sequence but also information about the degree of conservation that extends beyond the requirements of the underlying amino acid sequence in coding regions. This information is important for complementing the overall degree of sequence conservation, in particular in single-stranded regions of the conserved RNA secondary structures like apical loops or bulges. In the 5BSL2 stem-loop, we encountered examples of an incongruent mode of evolution, where sequence and structure are conserved, but the base pairs realizing the structure are not formed by homologous nucleotides. Such situations may be indicative of functionally independent selection pressures on sequence and structure [60]. As a consequence, part of the conserved structure is not detectable in a sequence-based alignment. Evolutionary incongruencies thus may reduce the sensitivity of consensus structure prediction methods. The local nature of the “shifts” between sequence and structure, on the other hand, still makes it possible to detect larger structured elements, such as the 5BSL2 stem-loop, which contains a sufficient subset of congruent base pairs formed by homologous nucleotides.

All conserved RNA secondary structure models have been added to the `Rfam` database in version 14.10 or later. The alignment will serve as a standard for future work on HCV.

## Data Availability

The alignments are available in the supplementary information (see Files F5 - F9): (1) the nucleotide alignment with additional annotations such as RNA secondary structures and genes (F5); (2) the protein alignment with gene annotation (F6); and (3) the nucleotide and protein alignment combined with additional annotations such as RNA secondary structures and genes (F7). The RNA secondary structure models are provided in the `Rfam` database [55, 56].

## Author contributions statement

S.T. performed the clustering and alignment construction and wrote the manuscript. K.L. implemented the clustering and initial alignment construction. N.O. integrated the RNA secondary structures in the `Rfam` database. B.S. integrated the RNA secondary structures in the `Rfam` database and revised the subsection about the `Rfam` models. P.F.S. wrote the subsection about incongruent evolution. M.N. evaluated the alignment and wrote the biological interpretation of the results. A.I.P. integrated the RNA secondary structures in the `Rfam` database. M.M. conceived the original research idea of full-genome alignment with RNA secondary structure annotation, designed the study, performed alignment construction and refinement, and wrote the manuscript. All authors read and approved the final version of the manuscript.

## References

1. World Health Organization. Global progress report on HIV, viral hepatitis and sexually transmitted infections, 2021. Accountability for the global health sector strategies 2016–2021: actions for impact. World Health Organization, 2021.

2. Qui-Lim Choo, George Kuo, Amy J. Weiner, Lacy R. Overby, Daniel W. Bradley, and Michael Houghton. Isolation of a cDNA cLone Derived from a Blood-Borne Non-A, Non-B Viral Hepatitis Genome. Science, 244(4902):359–362, 1989.

3. G. Kuo, Q.-L. Choo, H.J. Alter, G.L. Gitnick, A.G. Redeker, R.H. Purcell, T. Miyamura, J.L. Dienstag, M.J. Alter, C.E. Stevens, G. E. Tegtmeier, F. Bonino, M. Colombo, W.-S. Lee, C. Kuo, K. Berger, J. R. Shuster, L. R. Overby, D. W. Bradley, and M. Houghton. An Assay for Circulating Antibodies to a Major Etiologic Virus of Human Non-A, Non-B Hepatitis. Science, 244(4902):362–364, 1989.

4. A Takamizawa, C Mori, I Fuke, S Manabe, S Murakami, J Fujita, E Onishi, T Andoh, I Yoshida, and H Okayama. Structure and organization of the hepatitis C virus genome isolated from human carriers. Journal of Virology, 65(3):1105–1113, 1991.

5. Derrick Chu, Songyang Ren, Stacy Hu, Wei Gang Wang, Aparna Subramanian, Deisy Contreras, Vidhya Kanagavel, Eric Chung, Justine Ko, Ranjit Singh Amirtham Jacob Appadorai, Sanjeev Sinha, Ziba Jalali, David W. Hardy, Samuel W. French, and Vaithilingaraja Arumugaswami. Systematic Analysis of Enhancer and Critical cis -Acting RNA Elements in the Protein-Encoding Region of the Hepatitis C Virus Genome. Journal of Virology, 87(10):5678–5696, 2013.

6. David M. Mauger, Michael Golden, Daisuke Yamane, Sara Williford, Stanley M. Lemon, Darren P. Martin, and Kevin M. Weeks. Functionally conserved architecture of hepatitis C virus RNA genomes. Proceedings of the National Academy of Sciences, 112(12):3692–3697, 2015.

7. Markus Fricke, Nadia Dünnes, Margarita Zayas, Ralf Bartenschlager, Michael Niepmann, and Manja Marz. Conserved RNA secondary structures and long-range interactions in hepatitis C viruses. RNA, 21(7):1219–1232, 2015.

8. Nathan Pirakitikulr, Andrew Kohlway, Brett D. Lindenbach, and Anna M. Pyle. The Coding Region of the HCV Genome Contains a Network of Regulatory RNA Structures. Molecular Cell, 62(1):111–120, 2016.

9. E. A. Brown, H. Zhang, L. H. Ping, and S. M. Lemon. Secondary structure of the 5’ nontranslated regions of hepatitis C virus and pestivirus genomic RNAs. Nucleic Acids Research, 20(19):5041–5045, 1992.

10. T. Tanaka, N. Kato, M. J. Cho, and K. Shimotohno. A novel sequence found at the 3’ terminus of hepatitis C virus genome. Biochemical and Biophysical Research Communications, 215(2):744–749, 1995.

11. K J Blight and C M Rice. Secondary structure determination of the conserved 98-base sequence at the 3’ terminus of hepatitis C virus genome RNA. Journal of Virology, 71(10):7345–7352, 1997.

12. Michael Niepmann, Lyudmila A. Shalamova, Gesche K. Gerresheim, and Oliver Rossbach. Signals Involved in Regulation of Hepatitis C Virus RNA Genome Translation and Replication. Frontiers in Microbiology, 9:395, 2018.

13. Cristina Romero-López and Alfredo Berzal-Herranz. The Role of the RNA-RNA Interactome in the Hepatitis C Virus Life Cycle. International Journal of Molecular Sciences, 21(4):1479, 2020.

14. Philipp Schult, Hanna Roth, Rebecca L. Adams, Caroline Mas, Lionel Imbert, Christian Orlik, Alessia Ruggieri, Anna M. Pyle, and Volker Lohmann. microRNA-122 amplifies hepatitis C virus translation by shaping the structure of the internal ribosomal entry site. Nature Communications, 9(1):2613, 2018.

15. K Tsukiyama-Kohara, N Iizuka, M Kohara, and A Nomoto. Internal ribosome entry site within hepatitis C virus RNA. Journal of Virology, 66(3):1476–1483, 1992.

16. Catherine L. Jopling, Minkyung Yi, Alissa M. Lancaster, Stanley M. Lemon, and Peter Sarnow. Modulation of hepatitis C virus RNA abundance by a liver-specific MicroRNA. Science (New York, N.Y.), 309(5740):1577–1581, 2005.

17. Jura Inga Henke, Dagmar Goergen, Junfeng Zheng, Yutong Song, Christian G. Schüttler, Carmen Fehr, Christiane Jünemann, and Michael Niepmann. microRNA-122 stimulates translation of hepatitis C virus RNA. The EMBO journal, 27(24):3300–3310, 2008.

18. Tetsuro Shimakami, Daisuke Yamane, Rohit K. Jangra, Brian J. Kempf, Carolyn Spaniel, David J. Barton, and Stanley M. Lemon. Stabilization of hepatitis C virus RNA by an Ago2-miR-122 complex. Proceedings of the National Academy of Sciences of the United States of America, 109(3):941–946, 2012.

19. Shihyun You, Decherd D. Stump, Andrea D. Branch, and Charles M. Rice. A cis-acting replication element in the sequence encoding the NS5B RNA-dependent RNA polymerase is required for hepatitis C virus RNA replication. Journal of Virology, 78(3):1352–1366, 2004.

20. Haekyung Lee, Hyukwoo Shin, Eckard Wimmer, and Aniko V. Paul. cis-acting RNA signals in the NS5B C-terminal coding sequence of the hepatitis C virus genome. Journal of Virology, 78(20):10865–10877, 2004.

21. Donald B. Smith, Jens Bukh, Carla Kuiken, A. Scott Muerhoff, Charles M. Rice, Jack T. Stapleton, and Peter Simmonds. Expanded classification of hepatitis C virus into 7 genotypes and 67 subtypes: Updated criteria and genotype assignment web resource. Hepatology, 59(1):318–327, 2014.

22. Sergio M Borgia, Charlotte Hedskog, Bandita Parhy, Robert H Hyland, Luisa M Stamm, Diana M Brainard, Mani G Subramanian, John G McHutchison, Hongmei Mo, Evguenia Svarovskaia, and Stephen D Shafran. Identification of a Novel Hepatitis C Virus Genotype From Punjab, India: Expanding Classification of Hepatitis C Virus Into 8 Genotypes. The Journal of Infectious Diseases, 218(11):1722–1729, 2018.

23. BV-BRC. Introducing the Bacterial and Viral Bioinformatics Resource Center (BV-BRC): a resource combining PATRIC, IRD and ViPR. Nucleic Acids Research, 51(D1):D678–D689, 2023.

24. L. McInnes, J. Healy, and S. Astels. hdbscan: Hierarchical density based clustering. The Journal of Open Source Software, 2(11), 2017.

25. W. Li and A. Godzik. Cd-hit: a fast program for clustering and comparing large sets of protein or nucleotide sequences. Bioinformatics, 22(13):1658–1659, 2006.

26. L. Fu, B. Niu, Z. Zhu, S. Wu, and W. Li. CD-HIT: accelerated for clustering the next-generation sequencing data. Bioinformatics, 28(23):3150–3152, 2012.

27. M. Steinegger and J. Söding. MMseqs2 enables sensitive protein sequence searching for the analysis of massive data sets. Nature Biotechnology, 35(11):1026–1028, 2017.

28. C. Mercier, F. Boyer, A. Bonin, and E. Coissac. SUMATRA and SUMACLUST: fast and exact comparison and clustering of sequences. Programs and Abstracts of the SeqBio 2013 Workshop, 2013. Available online at: https://git.metabarcoding.org/obitools/sumaclust/wikis/home/.

29. T. Rognes, T. Flouri, B. Nichols, C. Quince, and F. Mahé. VSEARCH: a versatile open source tool for metagenomics. PeerJ, 4:e2584, 2016.

30. K. Lamkiewicz and M. Marz. ViralClust - Find representative viruses for your dataset. 202x (in preparation), www.github.com/klamkiew/viralclust/.

31. Eric W. Sayers, Evan E. Bolton, J. Rodney Brister, Kathi Canese, Jessica Chan, Donald C. Comeau, Ryan Connor, Kathryn Funk, Chris Kelly, Sunghwan Kim, Tom Madej, Aron Marchler-Bauer, Christopher Lanczycki, Stacy Lathrop, Zhiyong Lu, Francoise Thibaud-Nissen, Terence Murphy, Lon Phan, Yuri Skripchenko, Tony Tse, Jiyao Wang, Rebecca Williams, Barton W. Trawick, Kim D. Pruitt, and Stephen T. Sherry. Database resources of the national center for biotechnology information. Nucleic Acids Research, 50(D1):D20–D26, 2022.

32. K. Katoh, K. Misawa, K.-i. Kuma, and T. Miyata. MAFFT: a novel method for rapid multiple sequence alignment based on fast Fourier transform. Nucleic Acids Research, 30(14):3059–3066, 2002.

33. S. Will, K. Reiche, I. L. Hofacker, P. F. Stadler, and R. Backofen. Inferring Non-Coding RNA Families and Classes by Means of Genome-Scale Structure-Based Clustering. PLoS Comput Biol, 3(4):e65, 2007.

34. Guido Van Rossum and Fred L. Drake. Python 3 Reference Manual. CreateSpace, 2009.

35. K. Lamkiewicz and M. Marz. VeGETA - Viral GEnome sTructure Alignments. 202x (in preparation), https://github.com/klamkiew/vegeta.

36. Markus Fricke, Ruman Gerst, Bashar Ibrahim, Michael Niepmann, and Manja Marz. Global importance of RNA secondary structures in protein-coding sequences. Bioinformatics (Oxford, England), 35(4):579–583, 2019.

37. Andrew E. Firth and Ian Brierley. Non-canonical translation in RNA viruses. The Journal of General Virology, 93(Pt 7):1385–1409, 2012.

38. M. Nassal, M. Junker-Niepmann, and H. Schaller. Translational inactivation of RNA function: discrimination against a subset of genomic transcripts during HBV nucleocapsid assembly. Cell, 63(6):1357–1363, 1990.

39. Tsung-Cheng Chang, Akio Yamashita, Chyi-Ying A. Chen, Yukiko Yamashita, Wenmiao Zhu, Simon Durdan, Avak Kahvejian, Nahum Sonenberg, and Ann-Bin Shyu. UNR, a new partner of poly(A)-binding protein, plays a key role in translationally coupled mRNA turnover mediated by the c-fos major coding-region determinant. Genes & Development, 18(16):2010–2023, 2004.

40. M. A. Larkin, G. Blackshields, N. P. Brown, R. Chenna, P. A. McGettigan, H. McWilliam, F. Valentin, I. M. Wallace, A. Wilm, R. Lopez, J. D. Thompson, T. J. Gibson, and D. G. Higgins. Clustal W and Clustal X version 2.0. Bioinformatics (Oxford, England), 23(21):2947–2948, 2007.

41. F. Jeanmougin, J. D. Thompson, M. Gouy, D. G. Higgins, and T. J. Gibson. Multiple sequence alignment with Clustal X. Trends in Biochemical Sciences, 23(10):403–405, 1998.

42. Michele Clamp, James Cuff, Stephen M. Searle, and Geoffrey J. Barton. The Jalview Java alignment editor. Bioinformatics, 20(3):426–427, 2004.

43. Sam Griffiths-Jones. RALEE–RNA ALignment editor in Emacs. Bioinformatics (Oxford, England), 21(2):257–259, 2005.

44. Laura K. McMullan, Arash Grakoui, Matthew J. Evans, Kathleen Mihalik, Montserrat Puig, Andrea D. Branch, Stephen M. Feinstone, and Charles M. Rice. Evidence for a functional RNA element in the hepatitis C virus core gene. Proceedings of the National Academy of Sciences of the United States of America, 104(8):2879–2884, 2007.

45. Niki Vassilaki, Peter Friebe, Philipe Meuleman, Stephanie Kallis, Artur Kaul, Glaucia Paranhos-Baccala, Geert Leroux-Roels, Penelope Mavromara, and Ralf Bartenschlager. Role of the hepatitis C virus core+1 open reading frame and core cis-acting RNA elements in viral RNA translation and replication. Journal of Virology, 82(23):11503–11515, 2008.

46. Peter Friebe, Julien Boudet, Jean-Pierre Simorre, and Ralf Bartenschlager. Kissing-loop interaction in the 3’ end of the hepatitis C virus genome essential for RNA replication. Journal of Virology, 79(1):380–392, 2005.

47. Cristina Romero-López, Alicia Barroso-Deljesus, Ana García-Sacristán, Carlos Briones, and Alfredo Berzal-Herranz. End-to-end crosstalk within the hepatitis C virus genome mediates the conformational switch of the 3’X-tail region. Nucleic Acids Research, 42(1):567–582, 2014.

48. Jes⃺s Castillo-Martínez, Lixin Fan, Mateusz P. Szewczyk, Yun-Xing Wang, and José Gallego. The low-resolution structural models of hepatitis C virus RNA subdomain 5BSL3.2 and its distal complex with domain 3’X point to conserved regulatory mechanisms within the Flaviviridae family. Nucleic Acids Research, 50(4):2287–2301, 2022.

49. Jing Zhang, Osamu Yamada, Takashi Sakamoto, Hiroshi Yoshida, Hiromasa Araki, Takayuki Murata, and Kunitada Shimotohno. Inhibition of hepatitis C virus replication by pol III-directed overexpression of RNA decoys corresponding to stem-loop structures in the NS5B coding region. Virology, 342(2):276–285, 2005.

50. Gäel Cristofari, Roland Ivanyi-Nagy, Caroline Gabus, Steeve Boulant, Jean-Pierre Lavergne, François Penin, and Jean-Luc Darlix. The hepatitis C virus Core protein is a potent nucleic acid chaperone that directs dimerization of the viral (+) strand RNA in vitro. Nucleic Acids Research, 32(8):2623–2631, 2004.

51. Sumangala Shetty, Seungtaek Kim, Tetsuro Shimakami, Stanley M. Lemon, and Mihaela-Rita Mihailescu. Hepatitis C virus genomic RNA dimerization is mediated via a kissing complex intermediate. RNA (New York, N.Y.), 16(5):913–925, 2010.

52. Marylin Rheault, Sophie E. Cousineau, Danielle R. Fox, Quinn H. Abram, and Selena M. Sagan. Elucidating the distinct contributions of miR-122 in the HCV life cycle reveals insights into virion assembly. Nucleic Acids Research, 51(5):2447–2463, 2023.

53. Blake A. Sweeney, David Hoksza, Eric P. Nawrocki, Carlos Eduardo Ribas, Fábio Madeira, Jamie J. Cannone, Robin Gutell, Aparna Maddala, Caeden D. Meade, Loren Dean Williams, Anton S. Petrov, Patricia P. Chan, Todd M. Lowe, Robert D. Finn, and Anton I. Petrov. R2DT is a framework for predicting and visualising RNA secondary structure using templates. Nature Communications, 12(1):3494, 2021.

54. Jesús Castillo-Martínez, Tamara Ovejero, Cristina Romero-López, Isaías Sanmartín, Beatriz Berzal-Herranz, Elisa Oltra, Alfredo Berzal-Herranz, and José Gallego. Structure and function analysis of the essential 3’X domain of hepatitis C virus. RNA (New York, N.Y.), 26(2):186–198, 2020.

55. S. Griffiths-Jones, A. Bateman, M. Marshall, A. Khanna, and S. R. Eddy. Rfam: an RNA family database. Nucleic Acids Research, 31(1):439–441, 2003.

56. I. Kalvari, E. P. Nawrocki, N. Ontiveros-Palacios, J. Argasinska, K. Lamkiewicz, M. Marz, S. Griffiths-Jones, C. Toffano-Nioche, D. Gautheret, Z. Weinberg, E. Rivas, S. R. Eddy, R. D. Finn, A. Bateman, and A. I. Petrov. Rfam 14: expanded coverage of metagenomic, viral and microRNA families. Nucleic Acids Research, 49(D1):D192–D200, 2020.

57. Elena Rivas. RNA structure prediction using positive and negative evolutionary information. PLOS Computational Biology, 16(10):e1008387, 2020.

58. Maria Waldl, Sebastian Will, Michael T. Wolfinger, Ivo L. Hofacker, and Peter F. Stadler. Bi-alignments as Models of Incongruent Evolution of RNA Sequence and Secondary Structure. In Paolo Cazzaniga, Daniela Besozzi, Ivan Merelli, and Luca Manzoni, editors, Computational Intelligence Methods for Bioinformatics and Biostatistics, volume 12313, pages 159–170. Springer International Publishing, Cham, 2020. Series Title: Lecture Notes in Computer Science.

59. Peter F. Stadler and Sebastian Will. Bi-alignments with affine gaps costs. Algorithms for Molecular Biology, 17(1):10, 2022.

60. Katja Nowick, Maria Beatriz Walter Costa, Christian Höner Zu Siederdissen, and Peter F. Stadler. Selection Pressures on RNA Sequences and Structures. Evolutionary Bioinformatics Online, 15:1176934319871919, 2019.

61. Gesche K. Gerresheim, Carolin S. Hess, Lyudmila A. Shalamova, Markus Fricke, Manja Marz, Dmitri E. Andreev, Ivan N. Shatsky, and Michael Niepmann. Ribosome Pausing at Inefficient Codons at the End of the Replicase Coding Region Is Important for Hepatitis C Virus Genome Replication. International Journal of Molecular Sciences, 21(18):6955, 2020.

62. Bui Quang Minh, Heiko A Schmidt, Olga Chernomor, Dominik Schrempf, Michael D Woodhams, Arndt Von Haeseler, and Robert Lanfear. IQ-TREE 2: New Models and Efficient Methods for Phylogenetic Inference in the Genomic Era. Molecular Biology and Evolution, 37(5):1530–1534, 2020.

63. Cristina Romero-López, Alicia Barroso-delJesus, and Alfredo Berzal-Herranz. The chaperone-like activity of the hepatitis C virus IRES and CRE elements regulates genome dimerization. Scientific Reports, 7:43415, 2017.

64. Michael Niepmann and Gesche K. Gerresheim. Hepatitis C Virus Translation Regulation. International Journal of Molecular Sciences, 21(7):2328, 2020.

