## Supplementary Information for "Comprehensive Survey of Conserved RNA Secondary Structures in Full-Genome Alignment of Hepatitis C Virus"

### Files

- F1 - original data set
- F2 - pre-filtered data set
- F3 - phylogenetic tree in `Newick (treefile)` and `Nexus (nex)` format
- F4 - representative genomes in `Fasta (fasta)` format
- F5 - nucleotide alignment in `ClustalW (aln)`, `Fasta (fasta)`, and `Stockholm (stk)` format (latter format with additional annotations such as RNA secondary structures and genes)
- F6 - protein alignment in `ClustalW (aln)`, `Fasta (fasta)`, and `Stockholm (stk)` format
- F7 - nucleotide and protein alignment in `Stockholm (stk)` format (with additional annotations such as RNA secondary structures and genes)
- F8 - alignment of the 5' UTR (51 sequences) in `Stockholm (stk)`, `ClustalW (aln)`, and `Fasta (fasta)` format
- F9 - alignment of the 3' X-tail (11 sequences) in `Stockholm (stk)`, `ClustalW (aln)`, and `Fasta (fasta)` format

### Clustering

**Table S1.** Clustering results of the initial HCV data set using `ViralClust` [30]. #Seqs – Number of Sequences; #Clstr – Number of Cluster; minClstr – Smallest Cluster; maxClstr – Largest Cluster; avClstr – Average Cluster Size; medClstr – Median Cluster Size; #UnclstrSeqs – Number of Unclustered Sequences

| Algorithm | #Seqs | #Clstr | minClstr | maxClstr | avClstr | medClstr | #UnclstrSeqs |
|---|---|---|---|---|---|---|---|
| `HDBSCAN` | 2 549 | 36 | 7 | 583 | 67.42 | 20 | 121 |
| `cd-hit-est` | 2 549 | 105 | 2 | 580 | 22.32 | 3 | 205 |
| `sumaclust` | 2 549 | 77 | 2 | 464 | 23.43 | 3 | 131 |
| `MMseqs2` | 2 549 | 100 | 2 | 623 | 23.59 | 3 | 190 |
| `vclust` | 2 549 | 124 | 2 | 535 | 19.22 | 3 | 166 |

### Genome completeness of representative genomes

- Complete Genome: NC_009823.1, NC_004102.1, NC_009827.1, NC_030791.1, AB047639.1, JF735122.1, MG717928.1, AF169004.1, JF735124.1, MH427311.1, AJ851228.1, KM504115.1, KJ470619.1, KJ439768.1, MK139017.1, DQ278891.1, EU158186.1, KJ678751.1, FJ462437.1, KC248199.1, KJ439777.1, JX227965.1, AY878652.1, MG717925.1, KY348757.1, EU234061.2 , EU234065.2
- Complete CDS: NC_009824.1, NC_009825.1, NC_009826.1, NC_038882.1, KC197233.1, MN628597.1, KM043284.1, MH590700.1, AB677533.1, KC197230.1, KC844040.1, KY620874.1, KY620603.1, MK327987.1, AY232740.1, MW689975.1, MW690013.1, MW689965.1, MN977327.1, MK548369.1, MW689971.1, AY587845.1, LC435023.1, KX767023.1, MW689962.1, MW689980.1, MG878999.1, MN164860.1, MN164851.1, MN164872.1

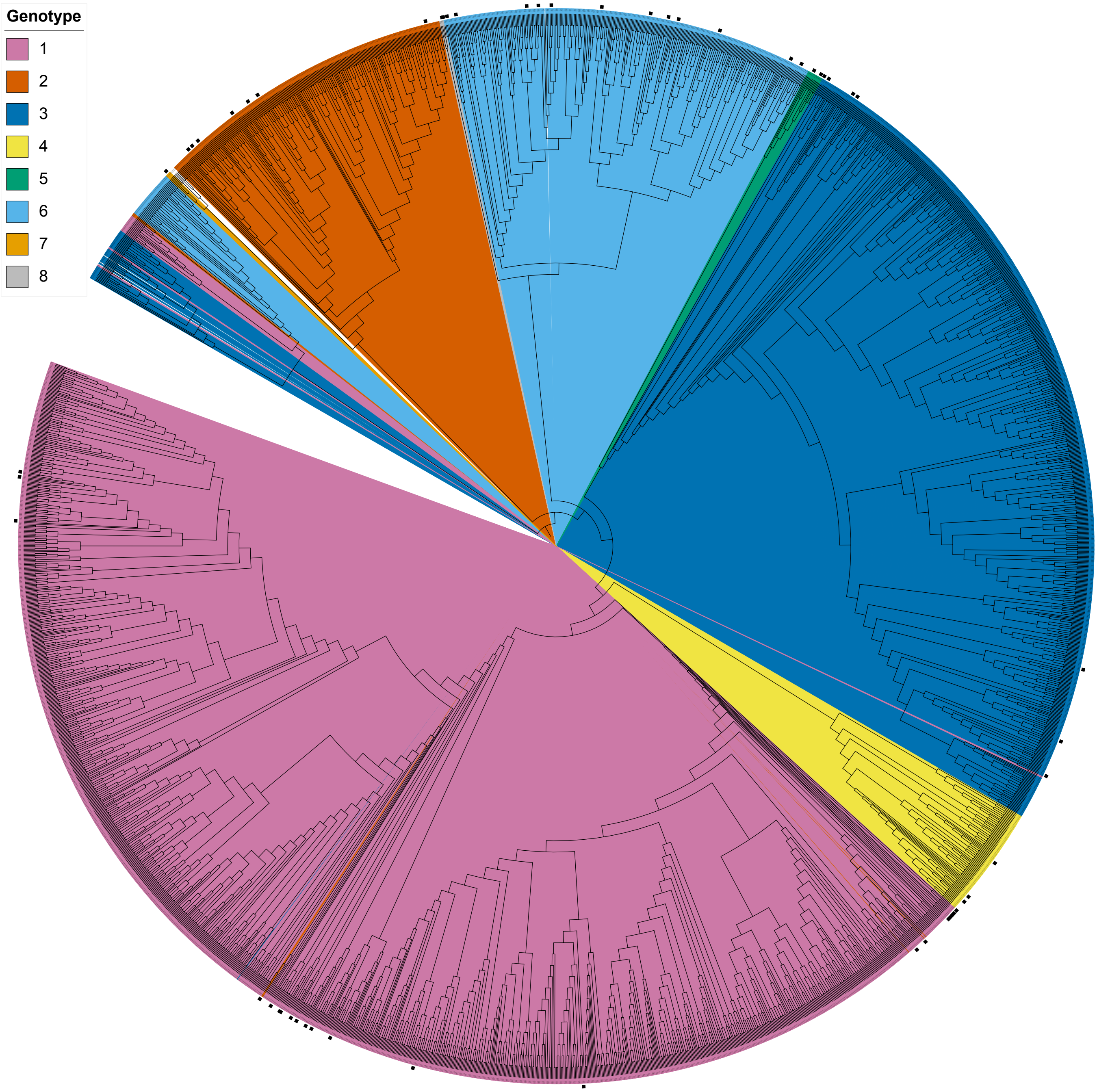


Figure S1: Phylogenetic tree of 2 549 HCV genomes obtained from `BV-BRC` database. The tree was reconstructed using `IQ-TREE 2` [62], employing an alignment generated by `MAFFT` [32]. The data set comprises all eight genotypes. While a few outliers are observed, the phylogenetic tree demonstrates clustering patterns corresponding to the different genotypes. The 57 representative genomes used for alignment construction are labeled with black boxes.

### Alignment confirms previously annotated RNA secondary structures – Additional Information

The full-genome alignment of HCV genomes reveals the presence of well-characterized RNA secondary structures, see Table 1, that are consistent with the existing literature [7, 8, 15]. These known structural elements include the highly conserved IRES (Figure S2), which plays a crucial role in HCV translation initiation [7, 15]. Additionally, stem-loop structures within the core region (Figure S5) and NS5B coding region (Figure S8 and Figure 1), such as the *cis*-replication element (CRE, composed of 5BSL3.1, 5BSL3.2, 5BSL3.3) and the highly conserved X-tail in the 3' UTR (both conformations) (Figure 2), were identified. Moreover, we predicted several conserved RNA secondary structures in the coding region of HCV genomes in agreement with the literature [7, 8]. The identification of these known structures not only validates the accuracy of our alignment but also underscores their functional importance in HCV biology: their conservation across different HCV genotypes further highlights their critical roles in viral replication, translation, and infection.

The importance of the highly conserved RNA secondary structures may be illustrated by a set of conserved functional RNA secondary structure elements in the C-terminal NS5B region in conjunction with 3' X sequences. The CRE/5BSL3.2 is important for HCV replication [19, 20]. Likely, in the early phase after HCV infection, the CRE apical loop can interact with the SL2 of the 3' X region by forming a "kissing loop" interaction [46], and this interaction may destabilize the dimerization linkage sequence (DLS) conformation of the upstream 3' X region but stabilize its SL2, SL3 conformation [47, 48]. In addition, a long-range interaction (LRI) between the CRE bulge and the classical form of the SLIIId of the IRES in the 5' UTR can form [7, 47, 63], likely facilitating translation [64]. In turn, at a time when sufficient amounts of NS5B replicase have been translated, NS5B can bind the CRE/5BSL3.2 and 5BSL3.3 [20, 49], thereby disabling the CRE – SL2 interaction and allow formation of the DLS [50, 51]. Binding of NS5B to the CRE then is supposed to be involved in starting RNA minus strand synthesis at the HCV RNA 3' end, whereas the role of the DLS and its putative role in dimerization of the full-length HCV genome in this process is not yet fully understood. These functional aspects are reflected by the high conservation of the involved RNA elements. The CRE/5BSL3.2 (Figure 1) is highly conserved in its secondary structure regions as well as in the apical loop and the bulge (in particular for the single-stranded regions, see also the conservation beyond coding sequence requirements, Figure S12). The same holds for the 5BSL3.3 which also binds NS5B [20]. The interacting RNA secondary structure, the SL2 of the 3' X region (Figure 2, left panel), is also highly conserved in both these aspects, see Figure 2 C). Moreover, the upstream 3' X region is conserved beyond the requirements of SL2 and 3 formation due to the overlapping constraints to form the DLS structure (Figure 2, right panel). These functional aspects in turn validate our alignment approach for identifying functionally important RNA secondary structures.

In the 5' and 3' UTRs, we would like to point to three cases in which alternative RNA secondary structure elements that both are functionally relevant may overlap and essentially share the same primary sequence. In general, this is a phenomenon that can be speculated to occur mainly in non-coding regions, since in coding regions the overlapping requirement for coding amino acids already applies strong constraints to the primary sequence, which may not allow for making more than one conserved RNA secondary structure. In the 5' UTR, these structures include 1) the SL II region, 2) the base of the SL III including the small SL IIId, and 3) in the 3' UTR the overlap of the predicted SLs 2 and 3 with the DLS (which was already discussed above in the context of the CRE-3' X interaction).

Confirming our previous results [7], we find the conserved capability of the IRES sequence to form a predicted alternative conformation of the SL IIId, the SL IIId*, at the base of the domain III of the IRES (Figure S2 A and B, and also see the alignment with the `StructConsensus` and `Consensus` outputs in Figure S3). Also in our RNA secondary structure alignment presented here, covering the sequence space of virtually all HCV isolates, the alternative SL IIId* structure is conserved in the predicted IRES structure with an MFE slightly lower than the IRES structure that comes with the classical SL IIId form. Even though the classical SL IIId has been repeatedly experimentally validated and shown to be functionally important, the structure presented here may have functional importance. This hypothesis is based on the lower MFE of this SL IIId* structure and on the following aspects. The sequence `GCGAAA` in the apical loop of the predicted alternative SL IIId* is virtually completely conserved, although this sequence is also partially single-stranded in the classical SL IIId form, a situation that likely would allow for possible sequence variation. Moreover, the five base pair stem carrying the apical `GCGAAA` loop of the SL IIId* is structurally conserved by covariations, although the classical SL IIId overlaps with that sequence, underlining the importance of the SL IIId*. Last but not least, the stem of the alternative SL IIId* appears to be slightly more stable by the number of base pairs that can be formed. We can only speculate if this alternative SL IIId* represents a structure that may be important in the IRES when not bound to ribosomes, likely after NS5B has bound the CRE/5BSL3.2 and by that interferes with the LRI between SL IIId and the bulge of the CRE, i.e. in the course of the switch from translation to replication.

Also, the region including SL II and the preceding sequence between SLs I and II is conserved in the primary sequence to a similarly high extent (please see exchanges in alignment in the STK file). This is again due to two alternative conformations that fulfill different tasks in the viral life cycle [52]. The conformation with the classical form of SL II is stabilized by two complexes of miR-122 with Argonaute (AGO) protein which binds between SL I and SL II. This classical SL II conformation has roles in HCV genome replication [16], promoting translation [17] and stabilization of the genome against nucleolytic degradation [18] during the initial phase of the infection, whereas the function promoting translation is rather dominant in late infection phases [52]. The alternative conformation SL II$^{alt}$ [14], however, appears to have a role in HCV assembly [52]. Therefore, this sequence region is highly conserved because it contains the two overlapping and alternating RNA secondary structures which both apply constraints to the primary sequence.

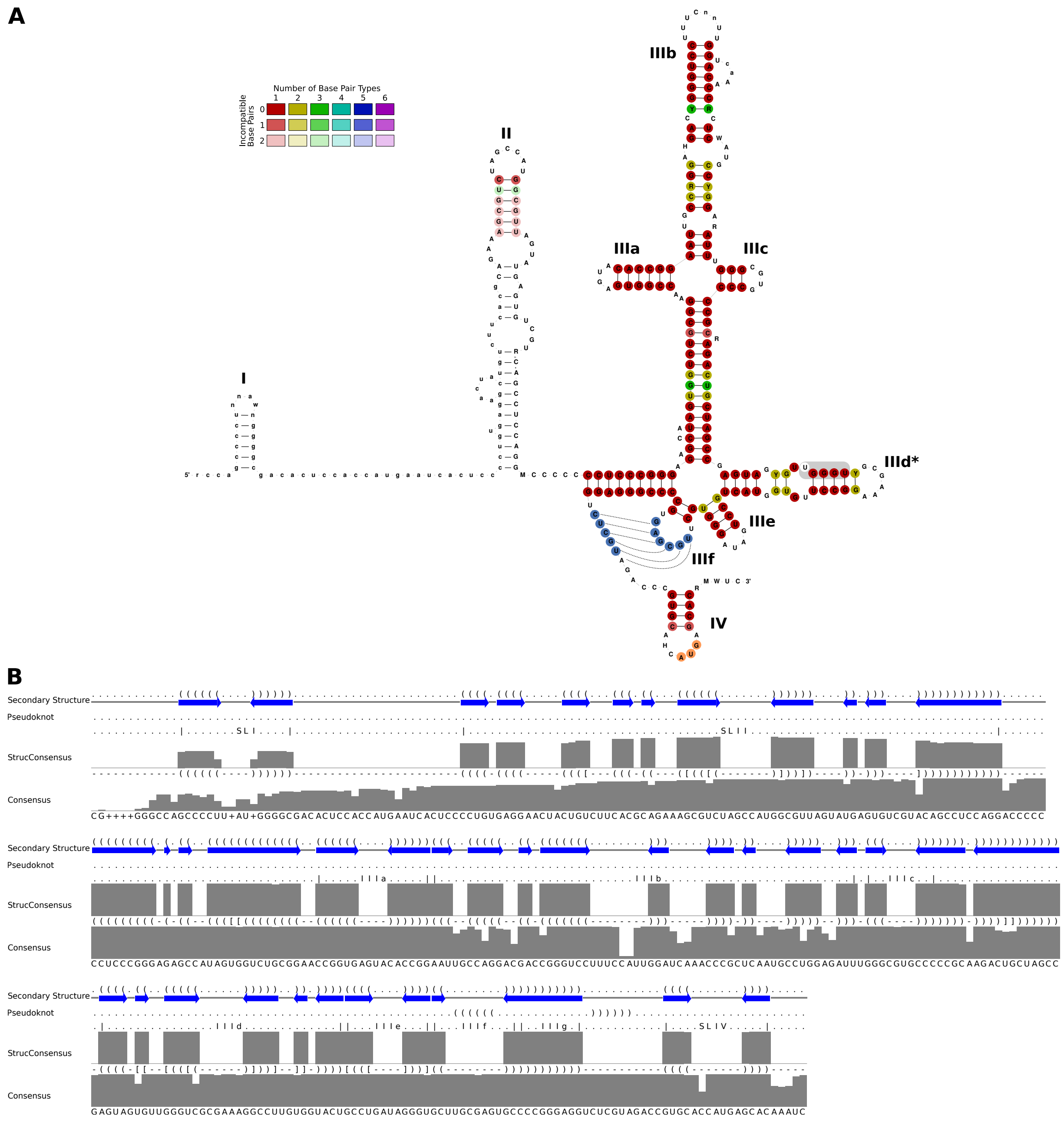


Figure S2: Conservation of the HCV 5' UTR. (A) Alignment-based RNA secondary structure prediction of the 5' UTR. Among the selected sequences, 51 isolates had a 5' UTR sequence (please see the additional alignment of only 5' sequences in Figure S3 and F8). The HCV IRES comprises the stem-loops (SLs) II to IV, with the polyprotein start codon located in the loop of SL IV. The IRES structure shows the same alternative structure of SL IIId* as in [7] due to the slightly lower MFE (-116.90 kcal/mol) compared to the "classical" experimentally validated structure. The sequence **UGGGU** that would be in the apical loop of the classical form of SL IIId is marked by the gray box. Only the nucleotides involving the loop of SL IIIf and part of the single-stranded stretch upstream of SL IV were manually manipulated to form a pseudoknot (blue), while the sequences involved in this pseudoknot actually were predicted to be single-stranded in the structure alignment. A low degree of covariance (see color code) suggests that not only the RNA secondary structure shown here is important but there may be also a requirement for overlapping *cis*-elements. The start codon (AUG) is marked in orange. The lower parts of the SL II appear largely white because on 41 sequences contain the stem-loop. (B) Sequence and RNA secondary structure features as in Figure 1 B.

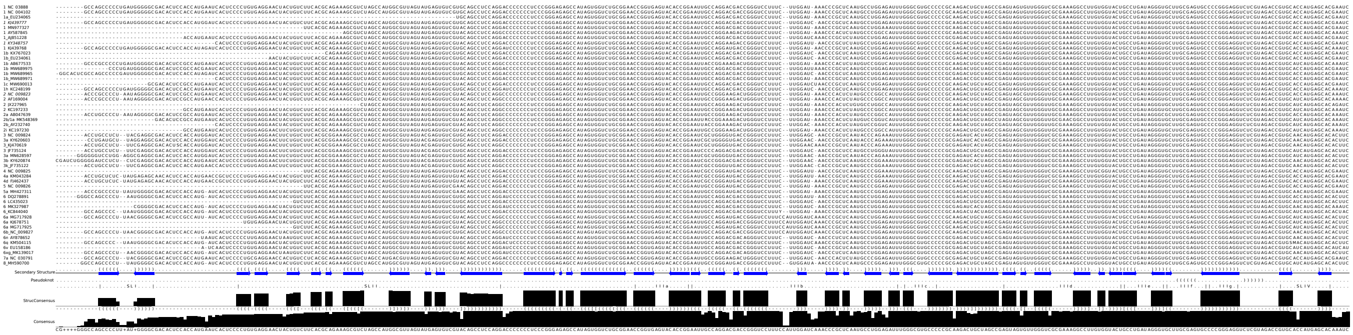

Figure S3: (related to Figure S2) Nucleotide alignment and additional sequence and RNA secondary structure features as in Figure 2 C of the 51 sequences fully covering the 5' UTR.

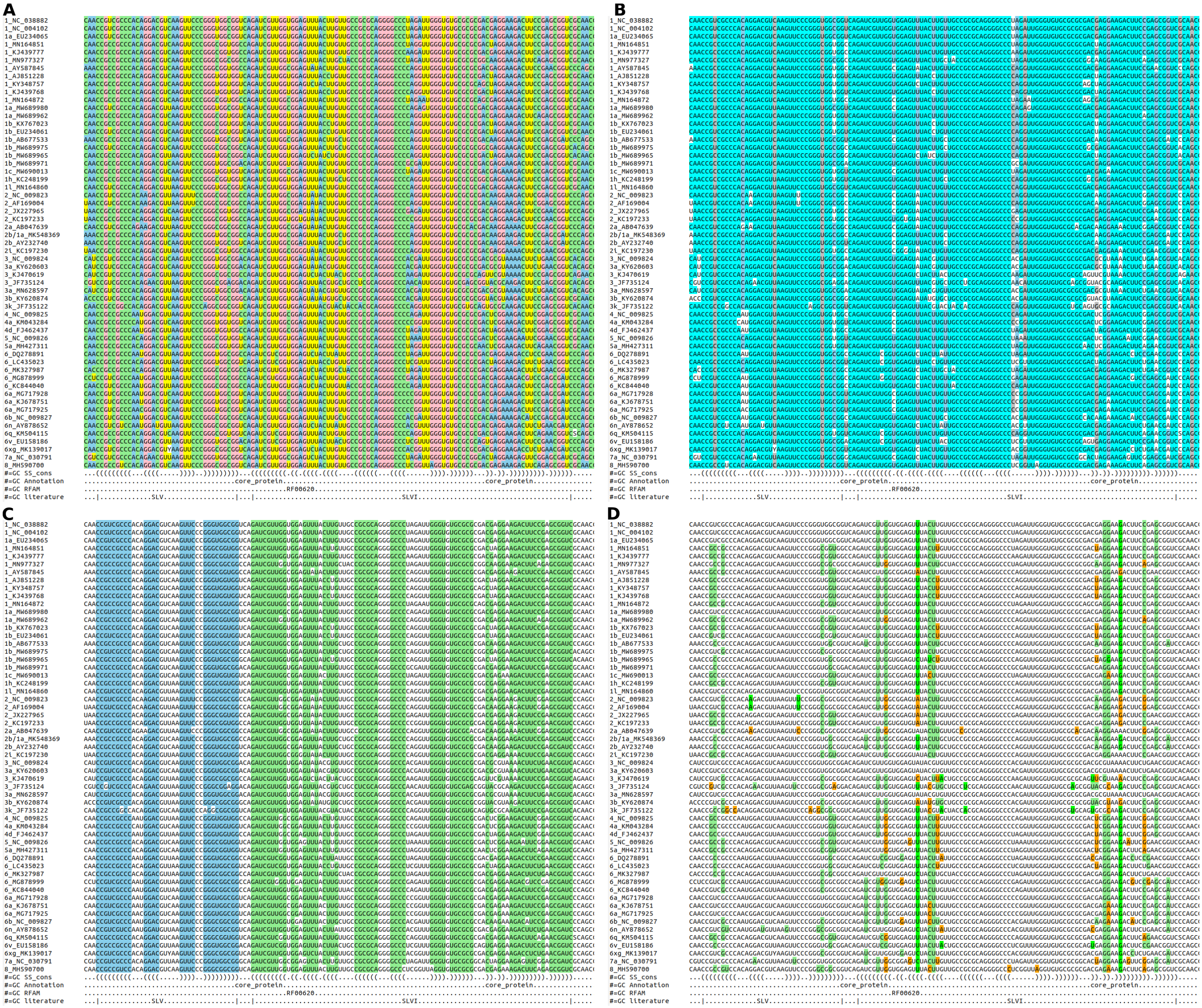

Figure S4: Nucleotide alignment visualization using **Emacs** in **RALEE** mode [43] displaying (A) nucleotide identity, (B) sequence conservation, (C) consensus RNA secondary structure, and (D) compensatory mutations.

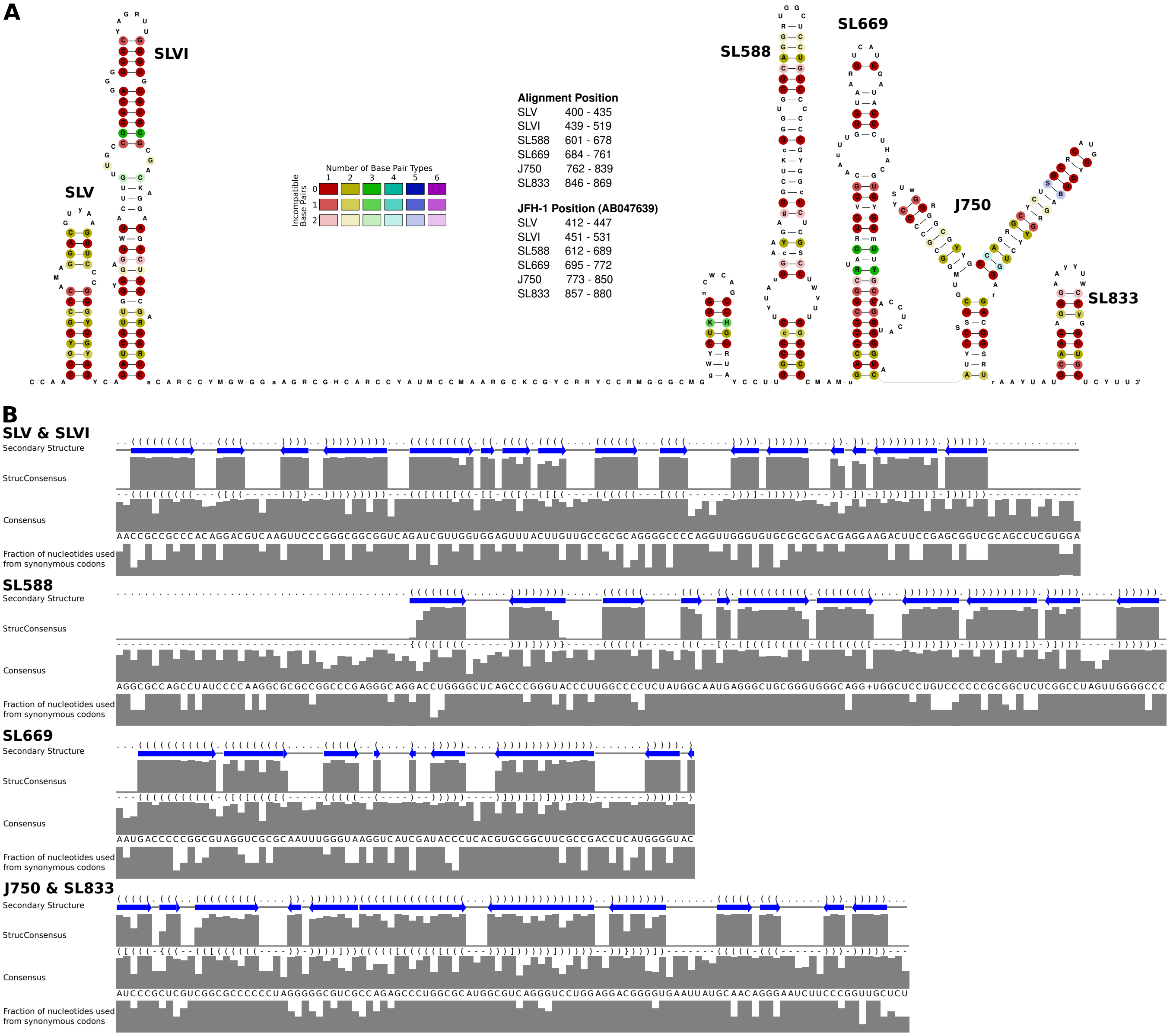


Figure S5: Conserved RNA secondary structures in the core coding region. (A) Structures shown with base color code illustrating the extent of covariations in double-stranded regions like in Figure S2. (B) Sequence and RNA secondary structure features as in Figure 1 B.

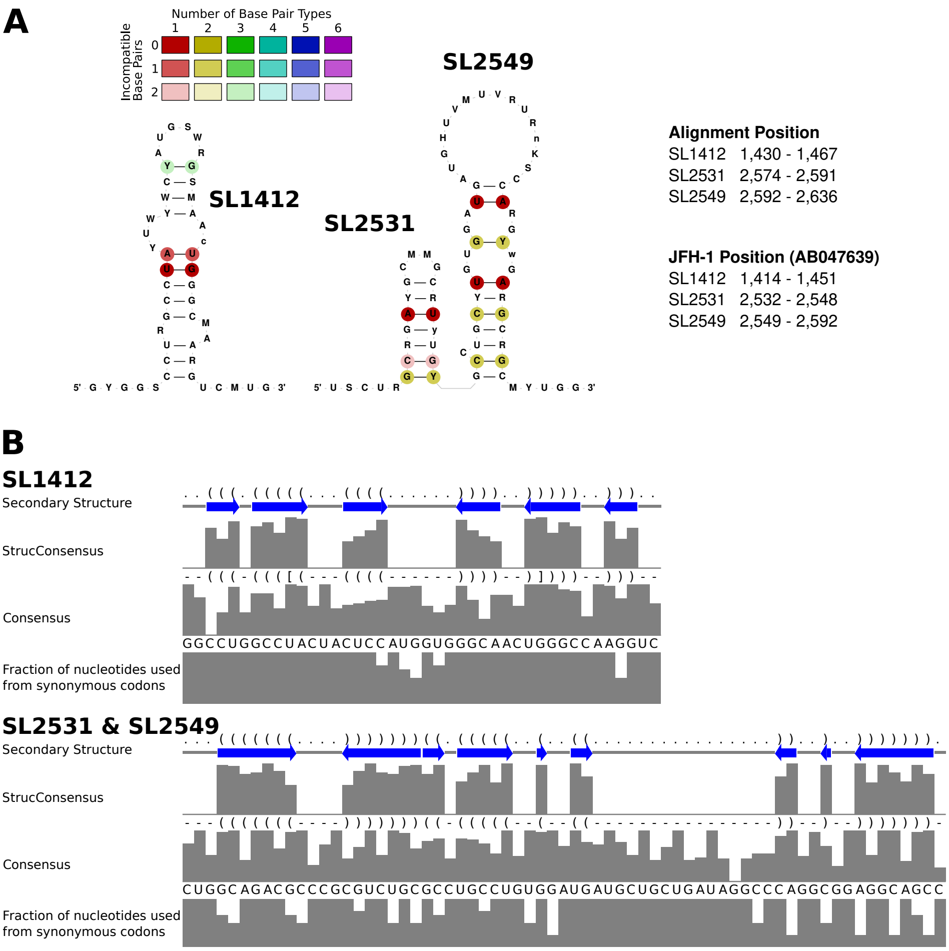


Figure S6: (A) Possible RNA secondary structures in the E1 and E2 region with its structural representation in and (B) additional sequence and RNA secondary structure features as in Figure 1 B.

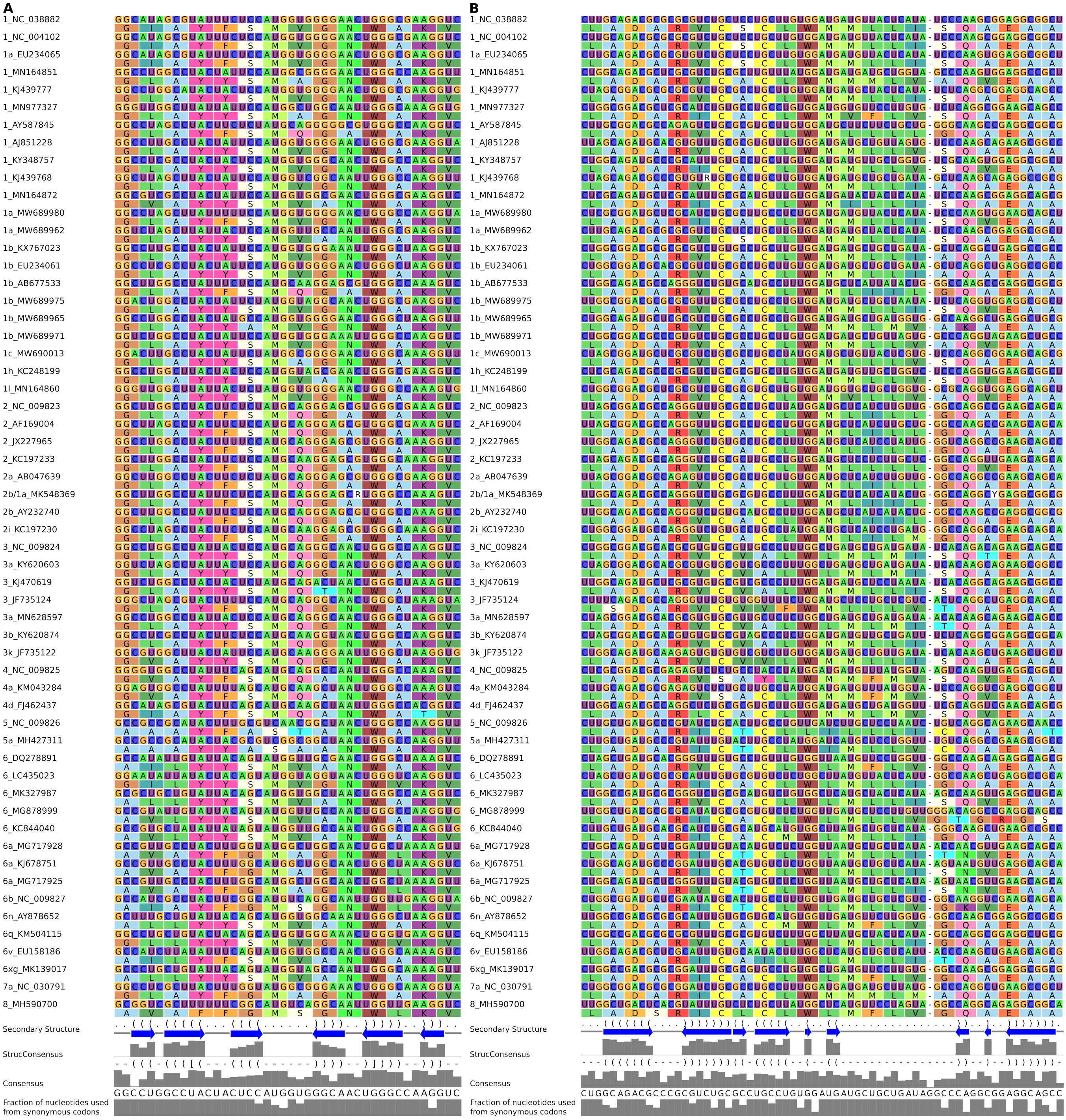


Figure S7: (related to Figure S6) Alignment of nucleotides and proteins combined and additional sequence and RNA secondary structure features as in Figure 1 B for (A) SL 1414 and (B) SL 2531 and SL 2549.

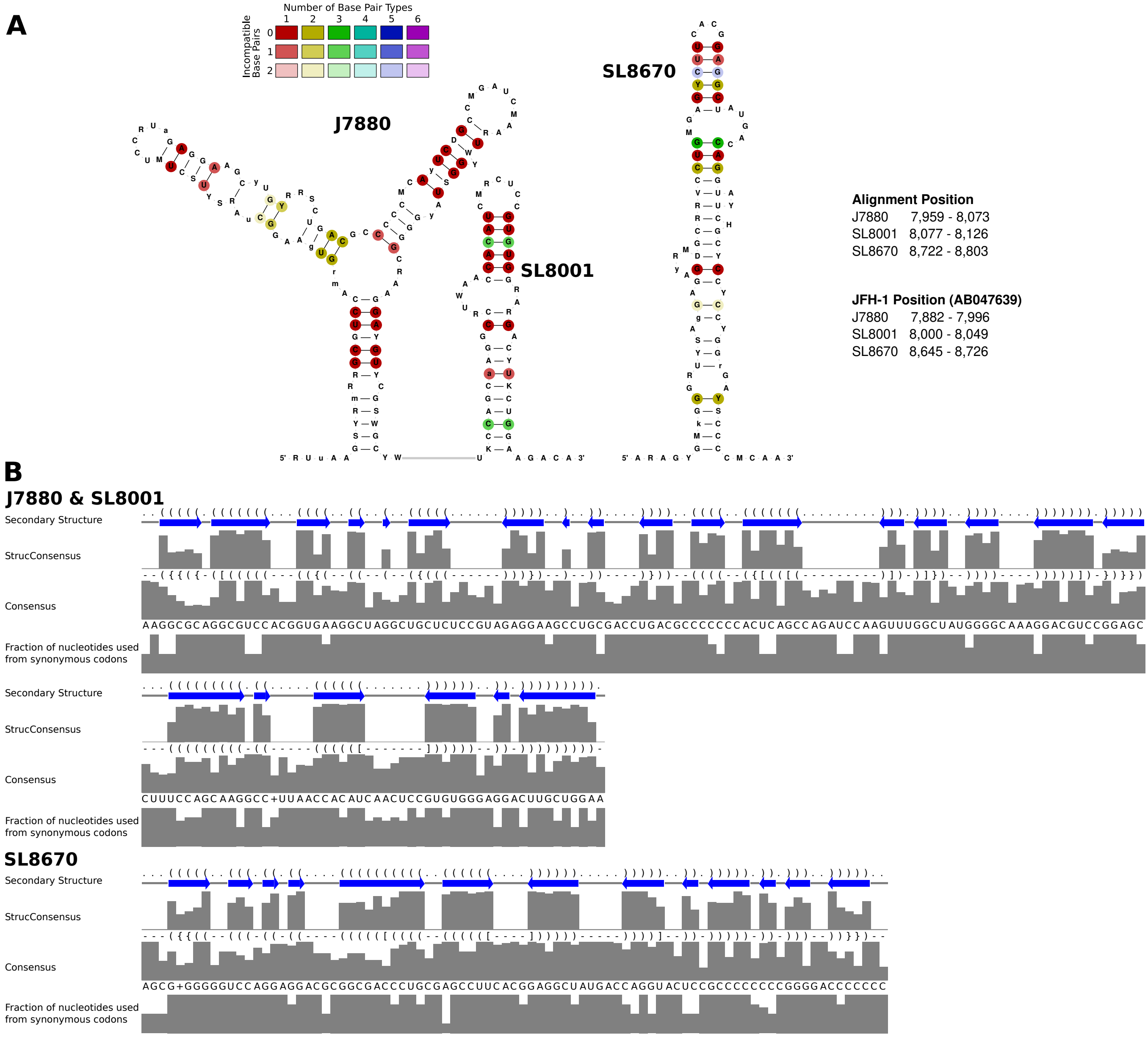


Figure S8: (A) Known RNA secondary structures in the upstream NS5B coding region and additional sequence and (B) RNA secondary structure features as in Figure 1 B.

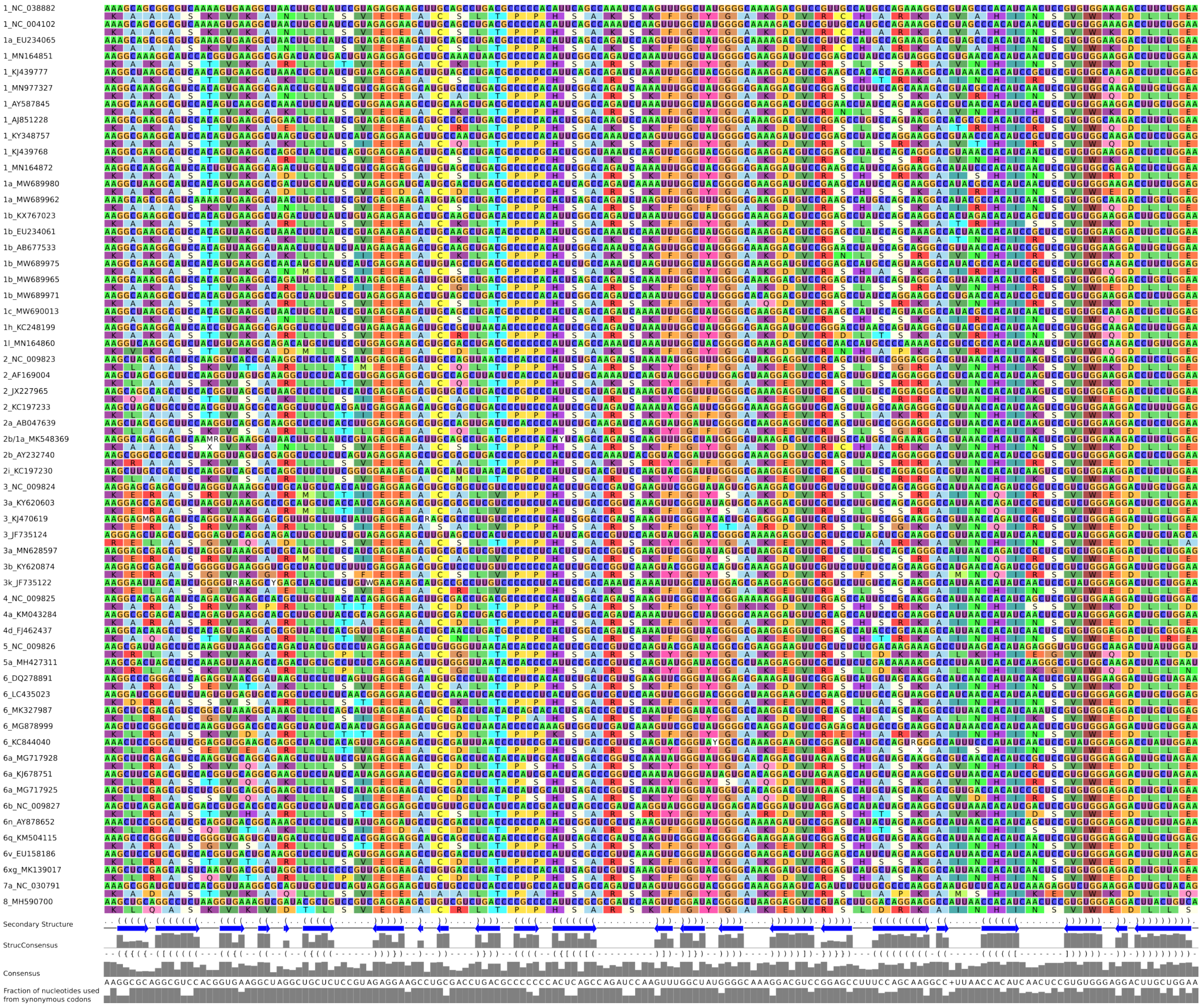

Figure S9: (related to Figure S8) Alignment of nucleotides and proteins combined and additional sequence and RNA secondary structure features as in Figure 1 B for J 7880 and SL 8001.

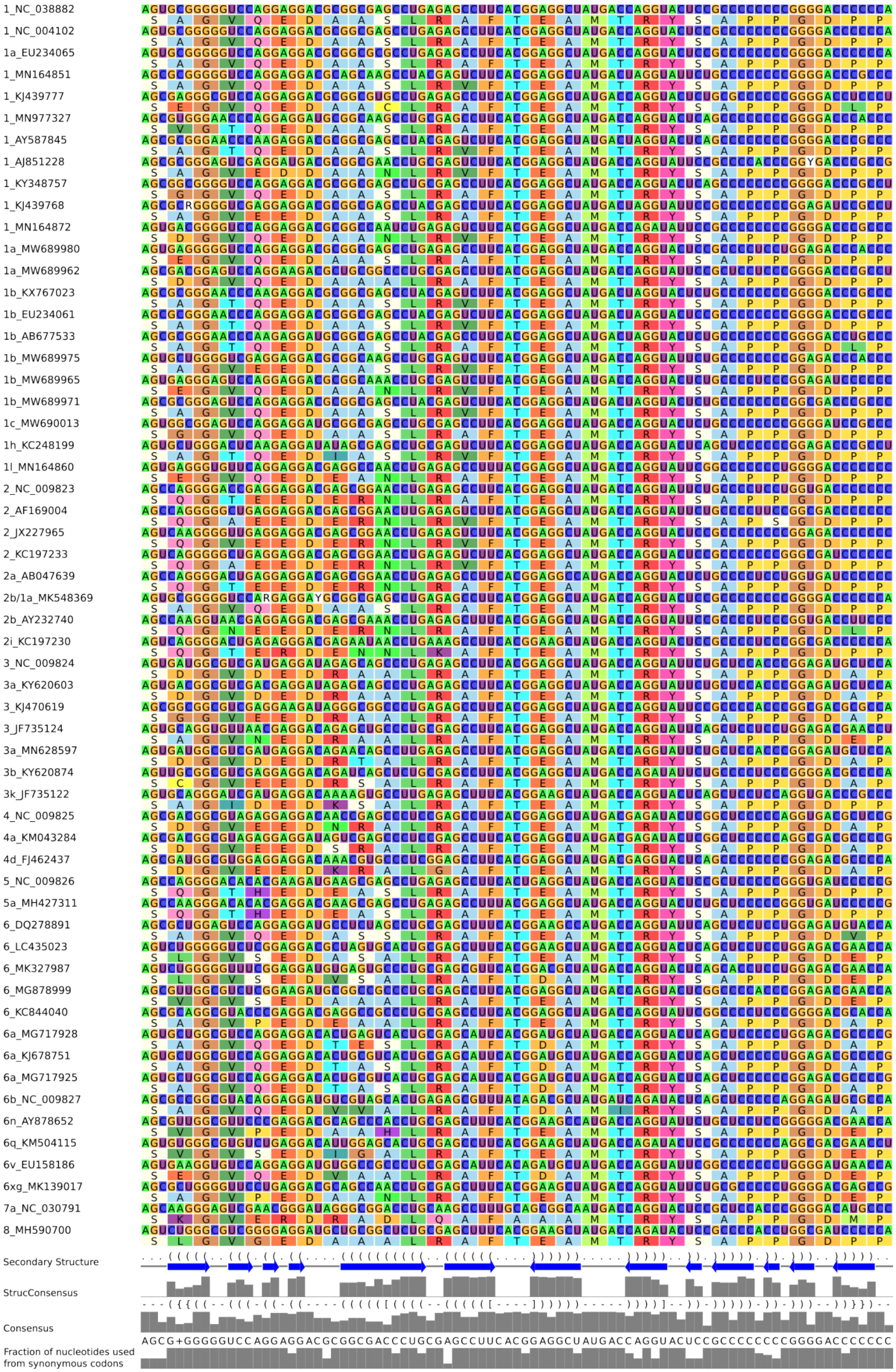


Figure S10: (related to Figure S8) Alignment of nucleotides and proteins combined and additional sequence and RNA secondary structure features as in Figure 1 B for SL 8670.

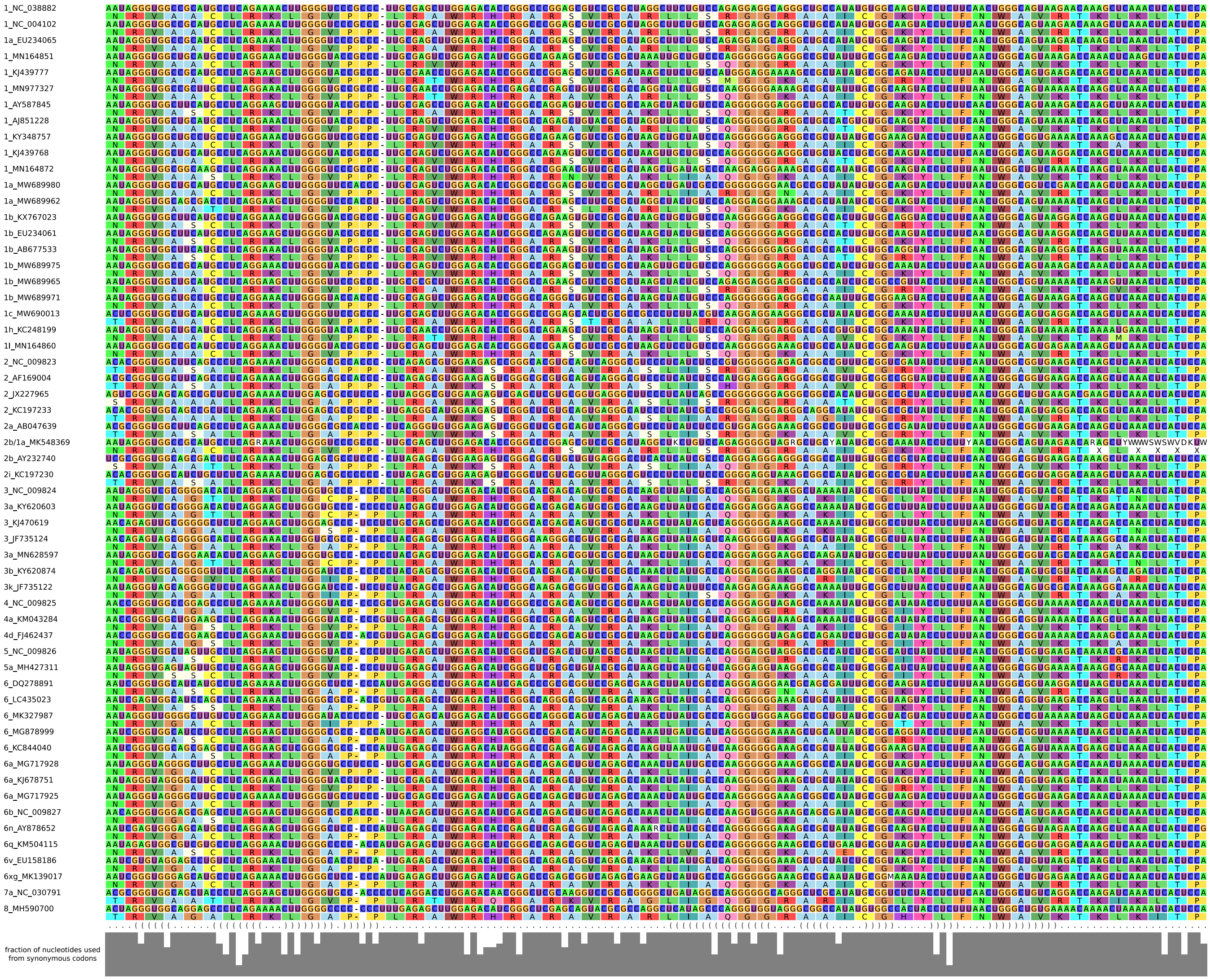

Figure S11: (related to Figure 1) Alignment of nucleotides and proteins combined and additional sequence and RNA secondary structure features as in Figure 1 B for 5BSL1 and 5BSL2.

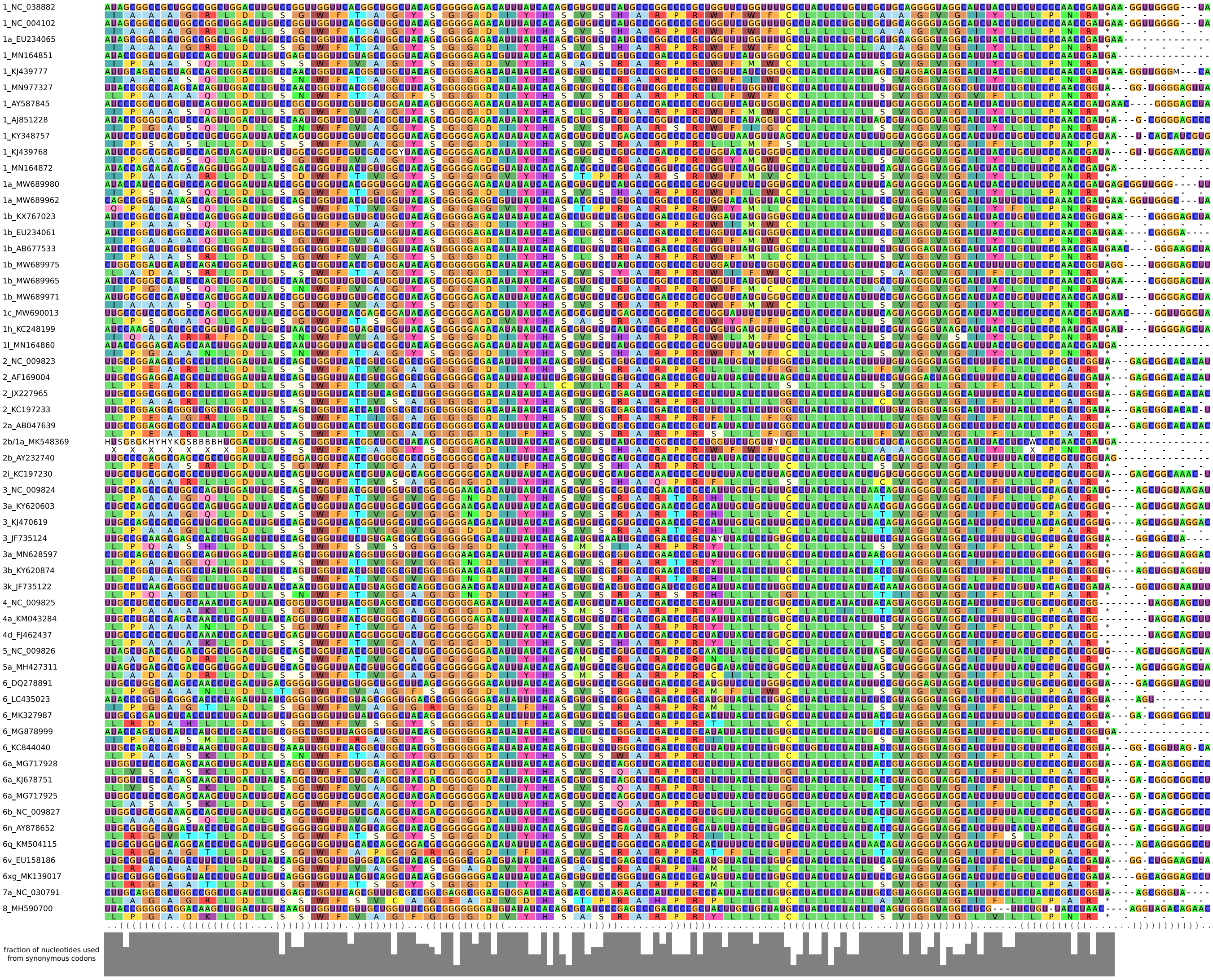


Figure S12: (related to Figure 1) Alignment of nucleotides and proteins combined and additional sequence and RNA secondary structure features as in Figure 1 B for *cis*-replication element (CRE, 5BSL3.2), 5BSL3.3 and SL IV.

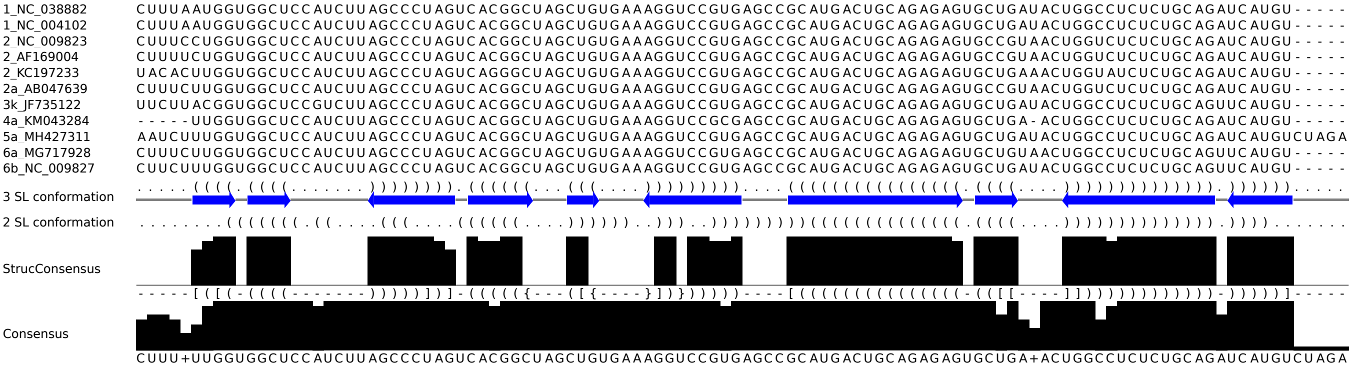


Figure S13: (related to Figure 2) Nucleotide alignment and additional sequence and RNA secondary structure features as in Figure 2 C of the 11 sequences fully covering the 3' X region.

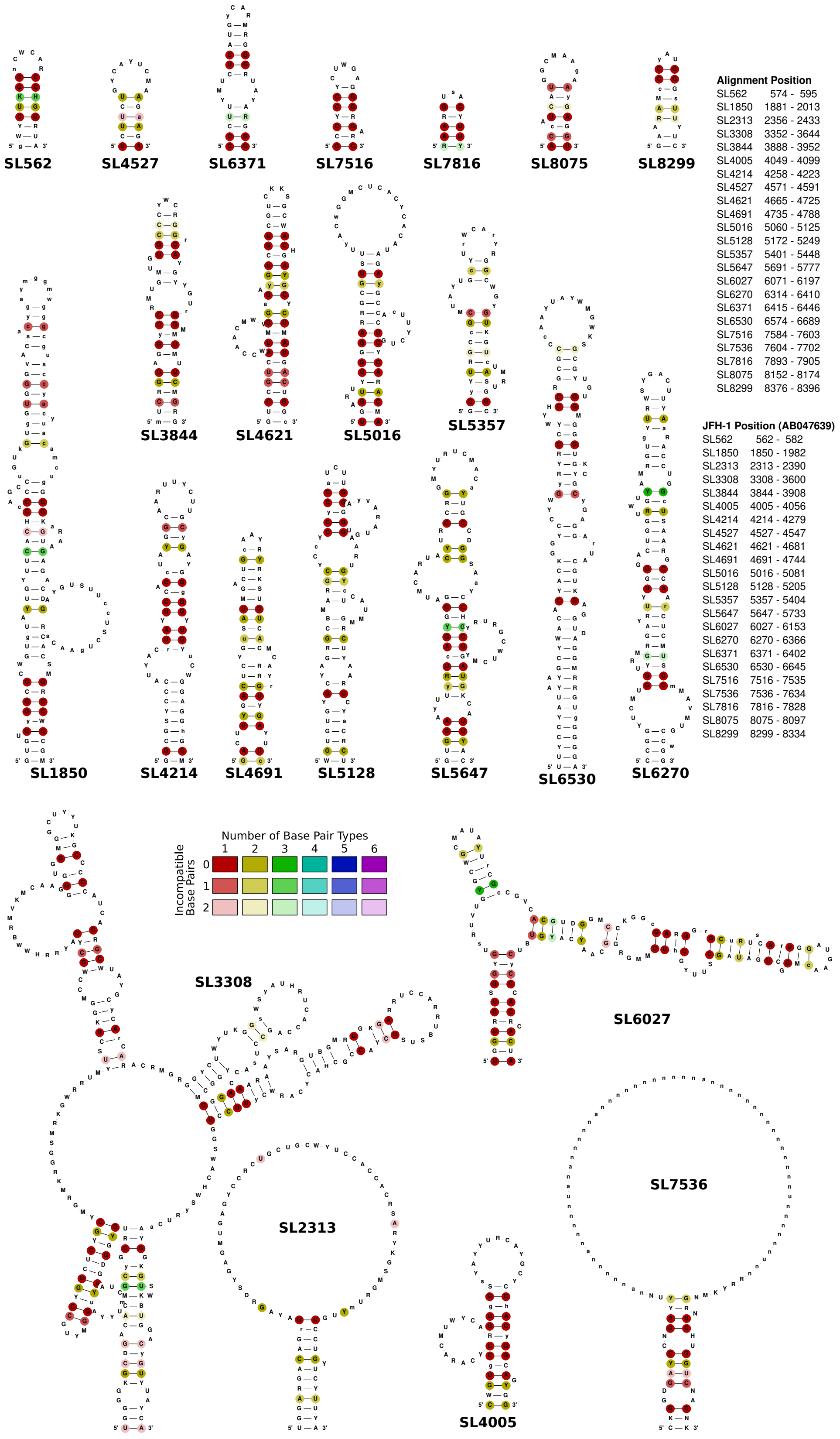


Figure S14: 23 novel conserved RNA secondary structure candidates from coding regions of the HCV alignment. RNA secondary structures are colored as in Figure 1 A.

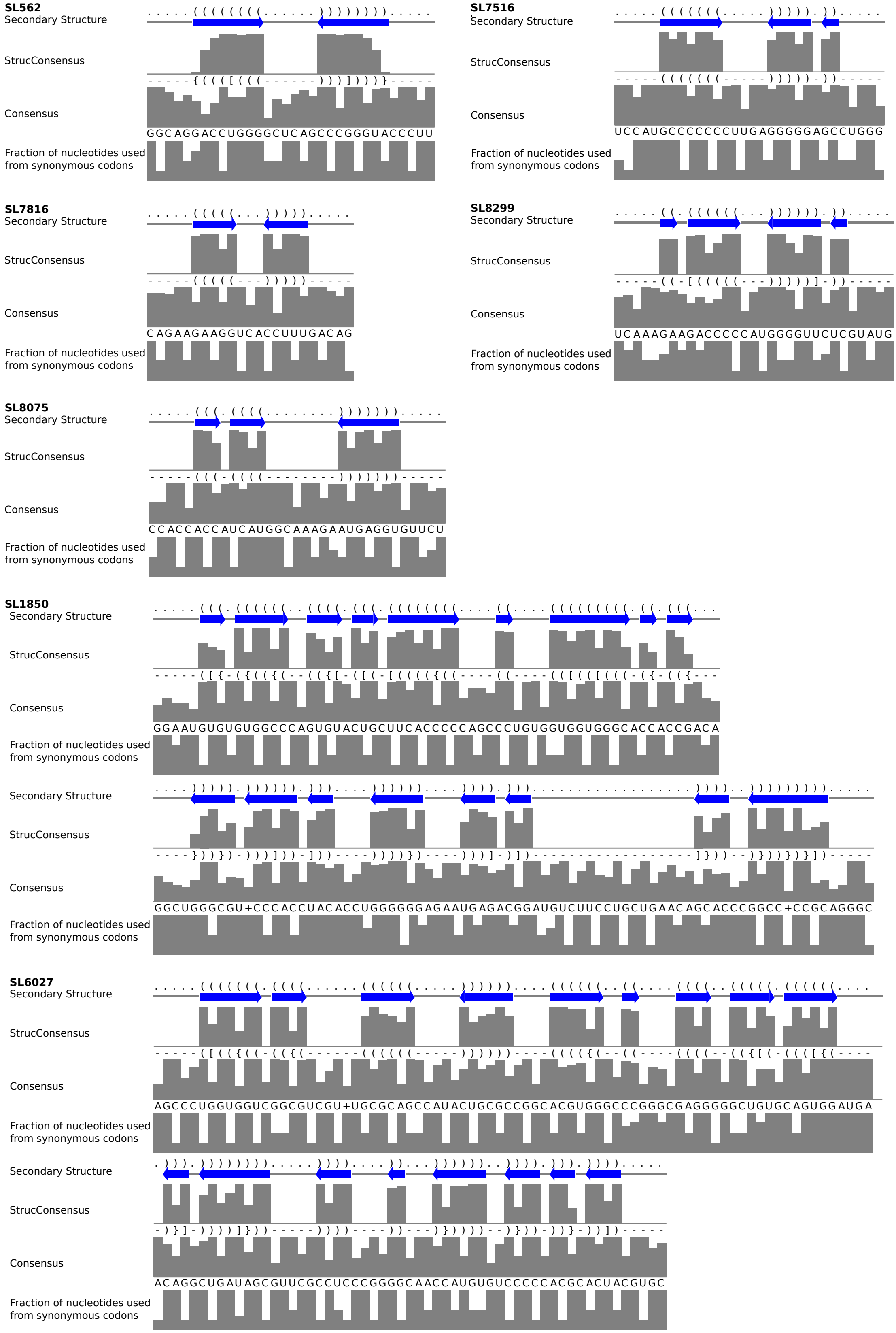


Figure S15: (related to Figure 3) Additional sequence and RNA secondary structure features as in Figure 1 B for seven novel conserved RNA secondary structure candidates.

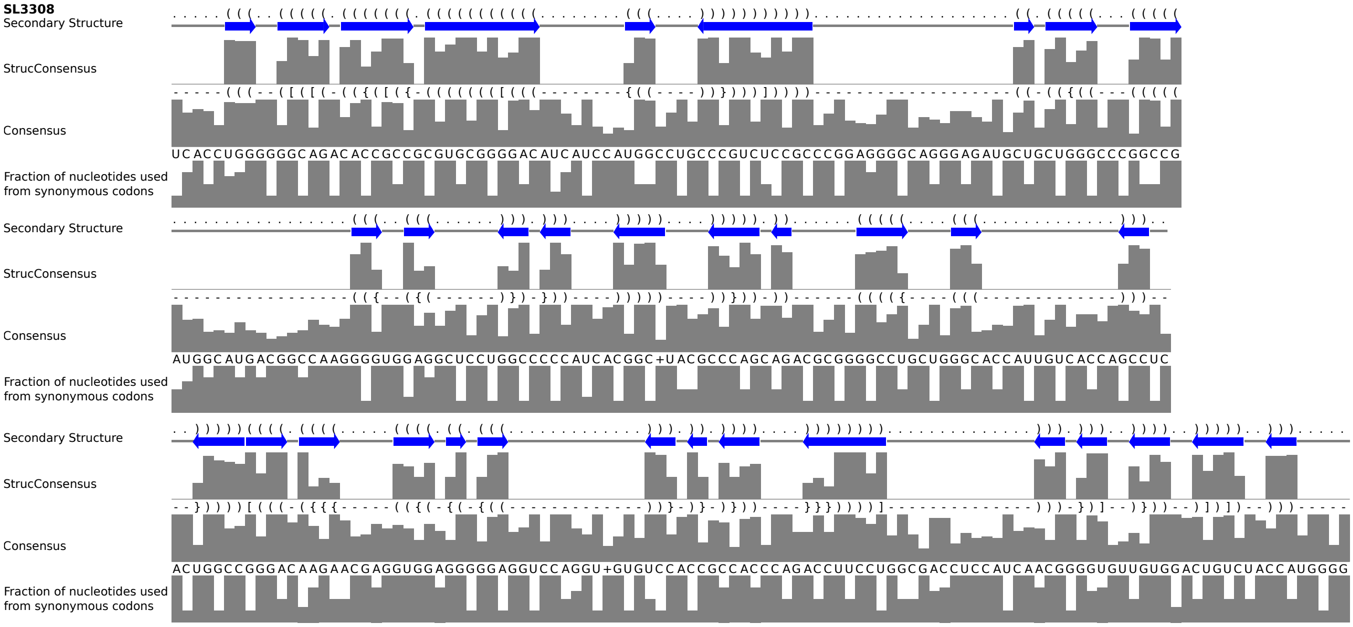


Figure S16: (related to Figure 3) Additional sequence and RNA secondary structure features as in Figure 1 B for SL 3308 as novel conserved RNA secondary structure candidate.
